# Human brain aging heterogeneity observed from multi-region omics data reveals a subtype closely related to Alzheimer’s disease

**DOI:** 10.1101/2024.03.01.582970

**Authors:** Shouneng Peng, Erming Wang, Minghui Wang, Xusheng Wang, Kaiwen Yu, Yingxue Fu, Suresh Poudel, Lap Ho, Sushma Narayan, Derek M. Huffman, Chris Gaiteri, David A. Bennet, Michelle E. Ehrlich, Vahram Haroutunian, Junmin Peng, Bin Zhang, Zhidong Tu

**Author notes:** Corresponding author: Zhidong Tu IMI 3-70F, 1425 Madison Ave New York City, NY 10029.

## Abstract

INTRODUCTION: The interconnection between brain aging and Alzheimer’s disease (AD) remain to be elucidated. METHODS: We investigated multi-omics (transcriptomics and proteomics) data from multiple brain regions (i.e., the hippocampus (HIPP), prefrontal cortex (PFC), and cerebellum (CRBL)) in cognitively normal individuals. RESULTS: We found that brain samples could be divided into ADL (AD-like) and NL (normal) subtypes which were correlated across brain regions. The differentially expressed genes in the ADL samples highly overlapped with AD gene signatures and the changes were consistent across brain regions (PFC and HIPP) in the multi-omics data. Intriguingly, the ADL subtype in PFC showed more differentially expressed genes than other brain regions, which could be explained by the baseline gene expression differences in the PFC NL samples. DISCUSSION: We conclude that brain aging heterogeneity widely exists, and our findings corroborate with the hypothesis that AD-related changes occur decades before the clinical manifestation of cognitive impairment in a sub-population.

## 1 Background

Brain aging is associated with cognitive decline, alterations in neuronal plasticity, dysregulation of proteostasis, accumulation of cellular damage, and elevation of inflammation[1]. These age-related changes are similar in some degree to those observed in mild cognitive impairment (MCI) and Alzheimer’s disease (AD), making it non-trivial to differentiate the aging-related cognitive decline from AD related changes in its early stages [2–5]. One approach to distinguish normative aging processes from those closely involved in AD development is to compare brain aging mechanisms in carefully selected samples: the aging mechanisms identified in an unimpaired brain aging should not be closely related to AD. By contrast, the aging mechanisms in individuals with “pathological aging” should be more closely related to AD. Here, we posit that brain aging is heterogeneous and can be generally divided into “normative” and “pathological” brain aging, each with distinct trajectories of cognitive performance in old age as supported by clinical data[6]. The key mechanisms of brain aging that drive the development of AD should be identifiable from the “pathological” brain aging subtype.

Understanding the molecular changes occurred in early stages of neurodegenerative diseases like AD may provide new insights into AD pathophysiology for drug target discovery, so that early interventions can be developed and introduced prior to the irreversible damages caused by neuropathology [7, 8]. Some evidence suggests that brain changes precedes the manifestation of dementia by about two or more decades [4, 5, 9]. For example, Beason-Held et al. showed that changes in brain function occurred years before the onset of cognitive impairment as indicated by changes in regional cerebral blood flow (rCBF)[10]. A brain imaging study covering N=4,329 subjects provides evidence of early divergence of the AD models from the normal aging trajectory before age 40 in the hippocampus[11]. A changepoint model studying 306 cognitively normal subjects with a subset developed AD in later years found that cerebrospinal fluid t-tau (CSF t-tau) had an estimated changepoint of approximately 34 years prior to symptom onset, although other markers were as short as 3 years[12]. All these studies suggest that we need to study donor’s human brain samples without cognitive impairment to understand the early molecular mechanisms that drive the development of AD.

Here, we hypothesize that brain aging is heterogenous and can be divided into subtypes; these subtypes can be identified or defined from brain tissue omics data; some of the subtypes may have undergone certain molecular changes that will lead to cognitive impairment even though the donors are cognitively normal at the time of death. To test this hypothesis, we leveraged multi-region (GTEx PFC and HIPP) and multi-omics (Resilience PFC transcriptomics and proteomics) datasets to identify brain aging subtypes using three different clustering methods: hierarchical clustering, K-means and weighted sample gene network analysis (WSCNA). It has been recognized that transcriptional profiling alone may not fully reveal the important modulation on proteins and higher-order cellular processes in brain aging and AD [13], therefore, the integration of transcriptomics and proteomics data could be helpful to unravel some unique mechanisms pertaining to brain aging heterogeneity [14]. Using this data-driven approach, we set the goal to better understanding brain aging heterogeneity at gene and protein levels among cognitively normal individuals, and to identifying the molecular link between brain aging heterogeneity and neurodegenerative diseases such as AD.

## 2 Materials and Methods

We collected multiple large-scale transcriptomics and proteomics datasets from cognitively unimpaired and AD brain samples to investigate brain aging heterogeneity. We describe each dataset below and summary information is listed in **Table S1**.

### 2.1 Genotype-Tissue Expression Brain Data

Genotype-tissue expression (GTEx) gene expression data (v8) from the prefrontal cortex (PFC), hippocampus (HIPP) and cerebellum (CRBL) were downloaded from the GTEx portal [15] and donors’ deidentified clinical information was obtained from the NIH dbGaP with access approval. Donors’ age ranged from 20 to 70 years. We further filtered out data from donors annotated with any brain/mental health related diseases from further analysis (i.e., MHALZDMT, MHALZHMR, MHCVD, MHDMNTIA, MHDPRSSN, MHENCEPHA, MHMENINA, MHPRKNSN, MHSCHZ). We adjusted sex, collection center (batch), RIN (RNA Integrity Number), PMI (postmortem interval), and top 3 genotype principal components (PCs) to obtain normalized gene expression. Low expressed genes with transcripts per every million reads sequenced (TPM) < 0.2 in more than 80% samples were removed.

### 2.2 Resilience (RS) data

RS data consists of 79 Mount Sinai Brain Bank (MSBB) samples from the PFC region and 88 ROSMAP samples from the dorsal lateral prefrontal cortical (DLPFC) region for a total of 167 normal aging samples, which have both gene expression and protein expression profiles available [16]. We used SMAP method [17] to align the transcriptomic and proteomic data to ensure samples from the two data types were accurately matched. Mis-aligned samples were removed from further joint omics analysis. Gene expression data was log2 transformed and adjusted for RIN, rRNA.rate, Intergenic.rate, Exonic.rate, batch, sex, race, PMI, sample. Protein expression data was log2 transformed and adjusted for batch, sex, race, PMI, sample. RS data is available for download from Synapse data portal (see Data Availability).

### 2.3 Seyfried2017 AD gene signatures

Seyfried2017 contained 50 individuals representing 15 controls, 15 AsymAD (Asymptomatic AD) and 20 AD cases from the Baltimore Longitudinal Study of Aging (BLSA)[18]. We only used data collected from the Brodmann Areas 9 (BA9) region.

### 2.4 Wingo2019 cognitive stability gene signatures

Wingo2019 contained discovery (NIJ=IJ104, 27% AD) and replication samples (NIJ=IJ39, 41% AD) of initially cognitively unimpaired, longitudinally assessed older-adult brain donors with Brodmann Areas 9 (BA9) samples. 2752 protein isoforms were detected in both Banner (Banner Sun Health Research Institute) and BLSA samples. 579 proteins were found to be associated with cognitive stability (350 up, 229 down) with consistent direction in both data after meta-analysis [19].

### 2.5 Mendonca2019 AD gene signatures

A total of 103 postmortem human brain samples from 46 subjects in 4 distinct brain regions (entorhinal cortex (BA28 and BA34), para-hippocampal cortex (posterior two thirds of the parahippocampal gyrus), temporal cortex (BA21) and frontal cortex (BA10)) were acquired from the Human Brain Tissue Bank (Semmelweis University, Budapest, Hungary). Analysis of differentially expressed proteins (DEPs) (adjusted p-value ≤ 0.05 and fold change ≤ 0.66 or ≥ 1.5) between the 4 brain regions in health state was performed using one-way ANOVA with post-hoc Tukey’s honestly significant difference (HSD) test [20]. We used data from the frontal cortex (BA10) in our analysis.

### 2.6 Ping2020 AD gene signatures

The dorsolateral prefrontal frontal cortex (BA9) samples were obtained from the Emory Alzheimer’s Disease Research Center (ADRC) brain bank. 27 samples from 3 groups (N = 10 control, N = 8 AsymAD, and N = 9 AD) were used for brain proteome analyses [21].

### 2.7 Jager AD gene signatures

Mostafavi et al. (2018) performed analyses on 478 ROSMAP dorsal lateral prefrontal cortex (DLPFC: BA9) tissue samples [22]. Five gene lists were considered which contained genes whose expression was associated with AD-related traits including clinical diagnosis of AD at the time of death, cognitive decline, tau, amyloid, and pathologic diagnosis of AD.

### 2.8 ROSMAP AD gene signatures

The Religious Orders Study and Memory and Aging Project (ROSMAP) dataset contained samples from dorsolateral prefrontal cortex (BA9) region (155 AD and 86 controls). DEGs were downloaded from the AMP-AD knowledge portal [23] and filtered for genes with FDR ≤ 0.05.

### 2.9 HBTRC BA9 AD gene signatures

Zhang et al. (2013) performed analyses on 549 postmortem specimens from BA9 in 376 LOAD and 173 nondemented subjects recruited through the Harvard Brain Tissue Resource Center (HBTRC). Each subject was diagnosed at intake and via extensive neuropathology examination [24]. Raw gene-expression data together with information related to demographics, disease state, and technical covariates are available via the GEO database GSE44772. DEG analyses were adjusted for age and sex, postmortem interval (PMI) in hours, and sample pH and RNA integrity number (RIN).

### 2.10 Canchi2019 AD gene signatures

This dataset contains gene expression profiles of 414 AD and non-demented controls from gray matter of BA9 brain region. After log-transformation and adjustment of covariates of age at death, sex, PMI and APOE status for the risk allele, DEGs were calculated using R package limma [25].

### 2.11 Annese2018 and Rooij2019 AD gene signatures

Annese2018 profiled hippocampus CA1 gene expression in 10 males with age between 60 ∼ 81(5 AD and 5 controls) [26]. Differentially expressed genes (DEGs) with | log2(Fold Change) | > 1 and FDR ≤ 0.05 were selected. Rooij2019 profiled gene expression of hippocampal samples from 18 AD and 10 controls. DEGs with differential expression score ≥ 0.1 and FDR ≤ 0.05 were considered [27].

### 2.12 AMP AD-PHG AD gene signatures

We obtained gene expression data from 215 parahippocampal gyrus (denoted as AMPAD_PHG) samples which were profiled at Mount Sinai [28]. To compare the PHG and GTEx hippocampal gene expression data, we first calculated log2(TPM + 1) for both datasets, merged these datasets and then removed the batch effects using R ComBat package with age, PMI, sex, RIN as covariates. We selected 78 samples [19 normal (CDR = 0, braak score: bbscore ≤ 3, CERAD = “NL”) and 59 LOAD (CDR ≥ 1, bbscore ≥ 5, CERAD = “definite AD”)] to define AD signature from this dataset using the same pipeline in our previous study [5].

### 2.13 Cognitive resilience signatures

Yu et al. identified 8 proteins ("NRN1", "ACTN4", "EPHX4", "RPH3A”/ “F8W131”/“RP3A”, "SGTB", "CPLX1", "SH3GL1", "UBA1”) which were associated with cognitive resilience (CR) based on dorsolateral prefrontal cortical data from 391 older adults (273 female, 118 male; mean ± SD age is 79.7 ± 6.7 years at baseline and 89.2 ± 6.5 years at death) in the ROSMAP study [29]. CR is the ability to maintain or improve cognitive function despite the presence, number, or combination of common brain pathologies, such as Alzheimer disease, Lewy bodies, transactive response DNA-binding protein 43, hippocampal sclerosis, infarcts, and vessel diseases. We further included two well-known cognitive resilient genes, *BDNF* [30] and *VGF* [31]. Among these ten genes, *UBA1* is the only down-regulated gene and the rest 9 are all up-regulated in the CR individuals.

### 2.14 Differential expression analysis and age-associated gene expression identification

Differentially expressed genes/proteins (DEGs/DEPs) analysis was performed using the R package edgeR and Limma [32, 33]. We adjusted batch, RIN, sex, and PMI in the GTEx data and adjusted RIN, rRNA.rate, intergenic.rate, exonic.rate, batch, sex, race, PMI, sample in the RS transcriptomics data and we adjusted batch, sex, race, PMI, sample in the RS proteomics data. We annotated the biological functions of each gene set using DAVID tool [34, 35].

### 2.15 Subgroup identification using hierarchical clustering, K-means and weighted sample gene network analysis (WSCNA)

We used hierarchical clustering, K-means and weighted sample gene network analysis (WSCNA) [36] to identify subgroups in GTEx and RS brain data. We selected the top 25,000 most variable genes for the hierarchical clustering of GTEx data and all genes in the RS data. Ward.D2 method in the R hclust function was used for the hierarchical clustering [37]. WSCNA calculates the similarity of samples by measuring the correlation of gene expression between a pair of samples. It also uses a topological overlap [topological overlap matrix (TOM)] score, which considers not only the correlation between two samples, but also the correlation of their shared neighbors in the network. To apply WSCNA, the gene expression data were normalized and adjusted for various confounding covariates. We used the default WSCNA settings and set beta to 6, RsquaredCut to 0.8 and minModuleSize to 5. We used clValid R package [38] to determine the optimal number of clusters for each dataset (see **Table S2**). We evaluated all possible models by setting K = 2 to 10 clusters and the choice for the number of clusters (K) was based on the stability and homogeneity of the clustering results and evaluated by two types of measures (internal and external) for the two classical clustering methods (Hierarchical, K-means), internal measures included the connectivity, Silhouette Width, and Dunn Index which were used to evaluate the connectedness, compactness and separation of the clusters. External measures were used to evaluate the stability of the clustering results included the average proportion of non-overlap (APN), and the average distance between means (ADM). These measures are particularly suitable for high-throughput genomic data, where the features often have high correlation [39].

### 2.16 Pseudo-temporal trajectory estimation

UMAP is a powerful tool for enhancing the diversity of samples in bulk -omic data and revealing meaningful clusters that correspond to biological and clinical features [40]. We followed the pseudo-temporal trajectory estimation procedure used by Wang et al. [41]. The adjusted expression data of GTEx (PFC, HIPP) was reduced to 50 dimensions based on a truncated principal component analysis (PCA) using an implicitly restarted Lanczos method in the R package Monocle3 [42]. The adjusted 59 genes (61 proteins) from RS (transcriptomics, proteomics) were not further dimensionality reduced. Data was further reduced to the first three dimensions using the Uniform Manifold Approximation and Projection (UMAP) [43] method from the R package uwot (https://github.com/jlmelville/uwot). We used a pseudotime index to measure the aging progression, namely the severity index (SI) along the trajectory. We calculated the SI (pseudotime) for each sample, based on the 3D UMAP trajectory, using the method of inferring pseudotime for single-cell transcriptomics from the function ‘slingPseudotime’ in the R package Slingshot [44].

### 2.17 Cross-brain region gene expression levels comparison

We compared the gene expression of NL/ADL samples between the PFC and HIPP to better understand region specific DEGs between NL and ADL samples. We put genes into several categories based on their expression levels in the NL and ADL samples across regions (FDR ≤ 0.01 was used to determine significant DEGs):

BaseNL_UP: UP-regulated genes in PFC ADL vs. NL, whose expression in PFC NL is significantly lower than its expression in HIPP NL, but their expression in PFC ADL is not significantly higher than its expression in HIPP ADL samples. These genes are PFC-specific DEGs mainly because their expression levels are lower in PFC NL samples (we refer to the gene expression level in PFC NL samples as the baseline expression).

BaseADL_UP: UP-regulated genes in PFC ADL vs. NL, whose expression in PFC ADL is significantly higher than its expression in HIPP ADL samples, but their expression in PFC NL is not significantly different from its expression in HIPP NL samples. These genes are PFC-specific DEGs mainly because their expression levels are substantially up-regulated in PFC ADL samples.

BaseALL_UP: UP-regulated genes in PFC ADL vs. NL, whose expression in PFC NL is significantly lower than its expression in HIPP NL and its expression is significantly higher in PFC ADL compared to HIPP ADL samples. These genes are PFC-specific DEGs mainly because their expression levels are lower in PFC NL samples and higher in PFC ADL samples.

BaseNL_DOWN, BaseADL_DOWN, and BaseALL_DOWN were similarly defined for the PFC specific down-regulated genes. The remaining DEGs were categorized as “Others DEGs” type.

### 2.18 Deconvolution of bulk tissue gene expression data to infer cell-type composition

We used the DSA (Digital Sorting Algorithm) [45] for cell-type proportion estimation [5, 46]. We followed our previous processing procedures (TMM normalization, top 100 markers) and applied DSA to the HIPP, PFC and PHG (parahippocampal gyrus) gene expression data (we adjusted age, sex, PMI, RIN, and batch) to estimate the cell-type proportions in these brain tissues.

## 3 Results

### 3.1 Datasets and AD gene/protein expression signatures considered in this study

We compiled a list of AD signatures from multiple large-scale human brain transcriptomics and proteomics datasets (**Tables S1A-C**). Briefly, AD transcriptomics signatures were obtained from HBTRC (BA9, 549 samples), Canchi2019 (BA9, 414 samples), ROSMAP (dorsolateral prefrontal cortex region (DLPFC/BA9, 241 samples), 5 lists from Jager’s AD gene signatures (DLPFC, 478 samples), AMP AD-PHG (PHG, 215 samples), Annese2018 (HIPP, 10 samples) and Rooij2019 (HIPP, 28 samples). AD proteomics signatures were obtained from Seyfried2017 (BA9, 50 samples), MendoncLa2019 (BA10, 26 samples), and Ping2020 (BA9, 27 samples). We also obtained cognitive trajectory proteomic signatures from Wingo2019 (BA9, 39 BLSA and 104 Banner samples). Our main focus is on analyzing cognitively normal brain data from the RS data (PFC transcriptomics and proteomics data) and the GTEx transcriptomics data in three brain regions (i.e., PFC, HIPP, and CRBL).

#### 3.1.1 Proteomic signatures of AD are negatively correlated with cognitive trajectory stability associated genes

To ensure that our brain aging and AD related proteomic signatures are of good quality and contain meaningful biological signals, we compared Wingo2019 cognitive stability signature with other AD proteomic signatures. As shown in **Figure S1A**, the cognitive stability of normative aging signatures (Wingo2019_BLSA, Wingo2019_Banner, Wingo2019_meta) showed reverse overlap with nearly every other AD proteomic signature, with the only exception for Ping2020_BA9_AsymNL AD signature. We reanalyzed the Ping2020 data and found that, as shown in **Figures S1B, S1C**, AD samples form a distinct group, except for two AD that mix with NL and AsymAD samples, while NL samples and AsymAD samples are more intermingled. Thus, the Ping2020_BA9_AsymNL list is not reliable. Based on this comparison, we removed Ping2020_BA9_AsymNL AD signatures and only used the remaining AD proteomic signatures for downstream analysis.

We defined a consistent AD proteomic signature by requiring that protein expression changes were present in at least 2 out of the 7 AD proteomic signature lists in the same direction, excluding the Ping2020_BA9_AsymNL AD signature. We denoted this consistent signature as the P_AD proteomic signature which consisted of 752 up-regulated genes and 1040 down-regulated genes. A comparison of the cognitive stability signature (Wingo2019 cognitive trajectory stability gene list from the meta-analysis) [19] with the P_AD proteomic signature suggests they are significantly overlapped in the opposite regulatory direction. For example, 239 PFC proteins are up-regulated in the cognitive stability signature but are down-regulated in the P_AD proteomic signature. These proteins are enriched for mitochondrial function, synapse, axon, dendritic spine, transport, energy metabolism (e.g., oxidative phosphorylation, respiratory chain, aerobic respiration, tricarboxylic acid cycle), post-translational modifications (PTMs) such as acetylation, and palmitoylation as annotated by the DVAID tool [34, 35] (see **Table S1D.1**). 113 PFC proteins are down-regulated in the cognitive stability signature but are up-regulated in the P_AD proteomic signature. These genes are enriched for extracellular exosome, PTMs (phosphoprotein, acetylation), cytoplasm, ficolin-1-rich granule lumen (3.4×10^-9^), and secretory granule lumen (8.2×10^-6^) (**Table S1D.2**).

#### 3.1.2 Brain prefrontal cortex AD transcriptomic and proteomic signatures highly overlap and are consistent in their regulatory directions

As shown in **Figure S1D**, except for the Ping2020_AsymNL AD signature, all AD transcriptomic lists (denoted as T_AD lists) and proteomic lists (P_AD lists) show significant overlap in the same regulatory direction (i.e., up-regulation in gene expression overlaps with up-regulation in protein expression and vice versa). We found that 1094 and 1068 genes (T_AD signature) were consistently up- and down-regulated in more than 2 gene lists in the 8 T_AD lists, respectively. We further overlapped the T_AD with the P_AD signature (752 up- and 1040 down-regulated proteins) which we named as the TP_AD signature. A total of 165 up-regulated and 258 down-regulated genes were contained in the TP_AD signature (**Figure S2**).

The function annotation of T_AD, P_AD and TP_AD gene signatures by the DAVID tool (with Benjamini-Hochberg adjusted FDR ≤ 0.05) [34, 35] is shown in **Figure S2** (detailed information is listed in **Table S1E**). Although there are fewer genes in the TP_AD signature compared to the T_AD signature, they have similar functions. For example, PTMs (phosphoprotein, acetylation), cytoplasm, focal adhesion, cytoskeleton and extracellular exosome are the top up-regulated functions; while synapse, axon, mitochondrion, transport, energy metabolism, membrane and ribosome related pathways are the top down-regulated functions. The P_AD-specific down-regulated signature is enriched for translation (FDR = 2.36E-04) and P_AD-specific up-regulated signature is enriched for transcription (FDR = 2.07E-12). T_AD specific down-regulated signature is enriched for rRNA processing (FDR = 1.54E-02) and T_AD specific up-regulated signature is enriched for mRNA transport (FDR = 1.88E-05) and mRNA splicing (FDR = 1.89E-05). This indicates that while the P_AD and T_AD signatures show significant common regulations, they also cover distinct aspects of molecular dysregulation, with the former affecting more on translation and transcription, while the latter affecting more on RNA processing and mRNA transport.

### 3.2 Multi-region clustering analysis shows two major subtypes in both the hippocampus and prefrontal cortex by a joint consideration of three clustering methods

Our previous study showed that healthy (NL) and AD-like (ADL) aging subtypes could be observed in both GTEx and UK hippocampal transcriptomics datasets [5]. Here, we extend this study to the GTEx PFC transcriptomic and RS PFC transcriptomics/proteomics data to test if brain aging subtypes can be observed in other brain regions (e.g., PFC and HIPP) and in different types of omics data (i.e., transcriptomics and proteomics data).

We applied the hierarchical and two other clustering methods, namely, K-means and WSCNA to define the subtype structures [36]. We used the R package clValid[38] to determine the optimal number of K-means and hierarchical clusters by considering the cluster number selection method proposed by Yang’s study [39]. We found that, for all the four datasets, the optimal number of clusters was 2 in most cases (see **Table S2**). For example, as shown in **Figure 1**, 129 GTEx hippocampal samples and 129 PFC samples (98 PFC/HIPP samples came from matched donors) showed two major subtypes based on hierarchical clustering (two main branches at the top level).

**Figure 1.**
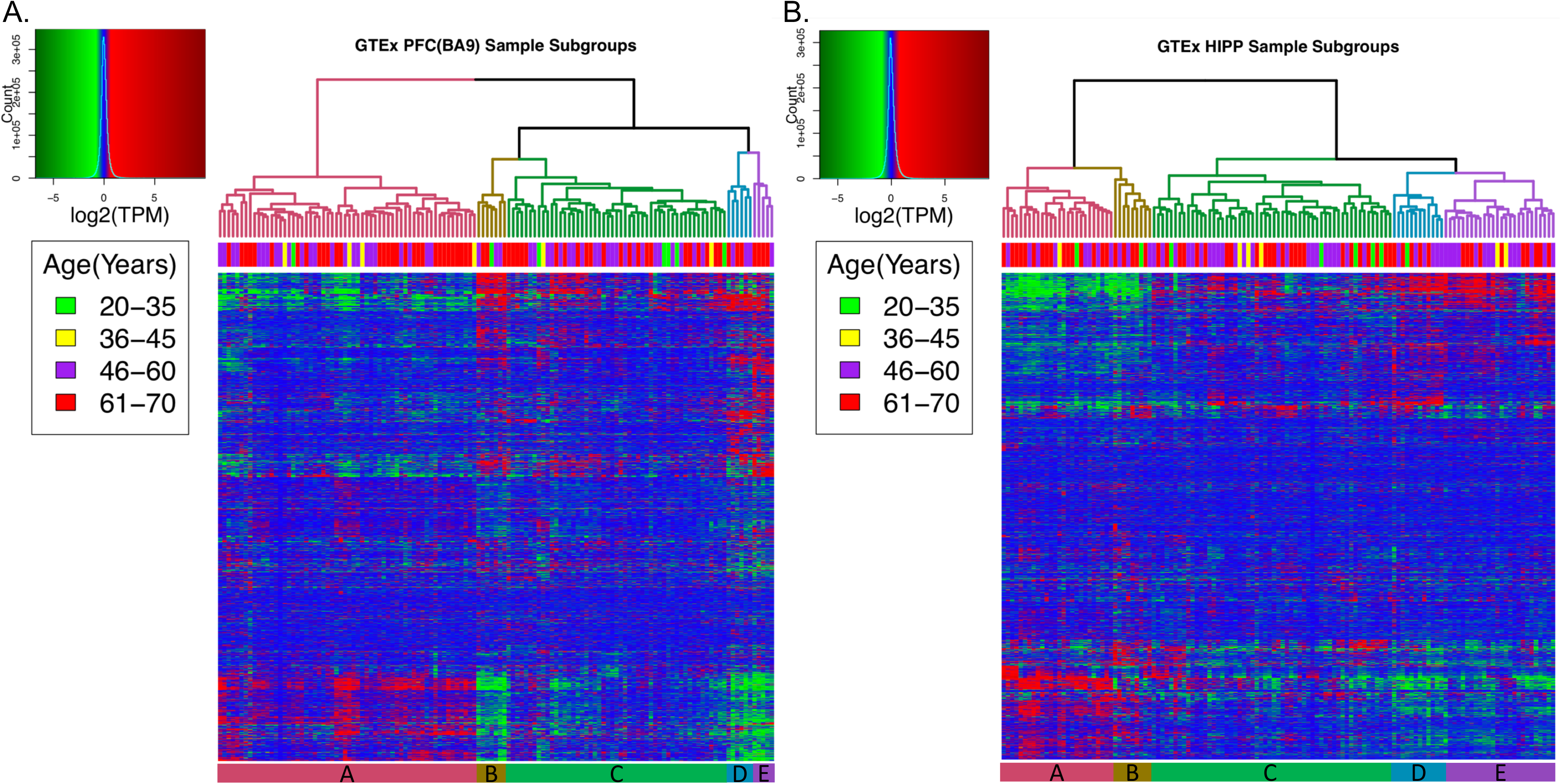
Multi-region (HIPP and PFC) clustering of brain transcriptomes showed two major subtypes which can be further divided into 5 subgroups (A, B, C, D, E, shown by different color in the dendrogram and labeled at the bottom of the heatmap) in each region. A. The subgroups of GTEx PFC; B. The subgroups of GTEx HIPP.

We then compared the three clustering methods considering two subtypes in each single dataset. Interestingly, we found that the two subtypes defined by each of the three clustering methods did not fully match with each other. As seen **Figure 2A**, when considering the top level two clusters, hierarchical cluster 1 (left side branch) could be perfectly mapped to K-means cluster 1 and WSCNA turquoise module. Hierarchical cluster 2 (right side branch) contained samples from both K-means clusters 1 and 2 (red and blue samples in the color bar) and did not match perfectly to either K-means or the WSCNA clusters. However, if we considered more subgroups (≥ 5) in the hierarchical clustering, we could better resolve the inconsistency. For example, by considering 5 PFC subgroups (A to E), we found that clusters B, D and E could now be well mapped to K-means cluster 2 (which is the ADL subtype), while A and C could be mapped to the K-means cluster 1 (which is the NL subtype) although the mapping of C was still imperfect. Since 5 clusters also received relatively high ranks by most methods implemented in clValid (**Table S2**) and having a larger number of subgroups would allow us to gain a higher resolution view of the specific cell types in each subgroup [5], we decided to consider 2 major subtypes which were further divided into 5 subgroups in our analysis.

**Figure 2.**
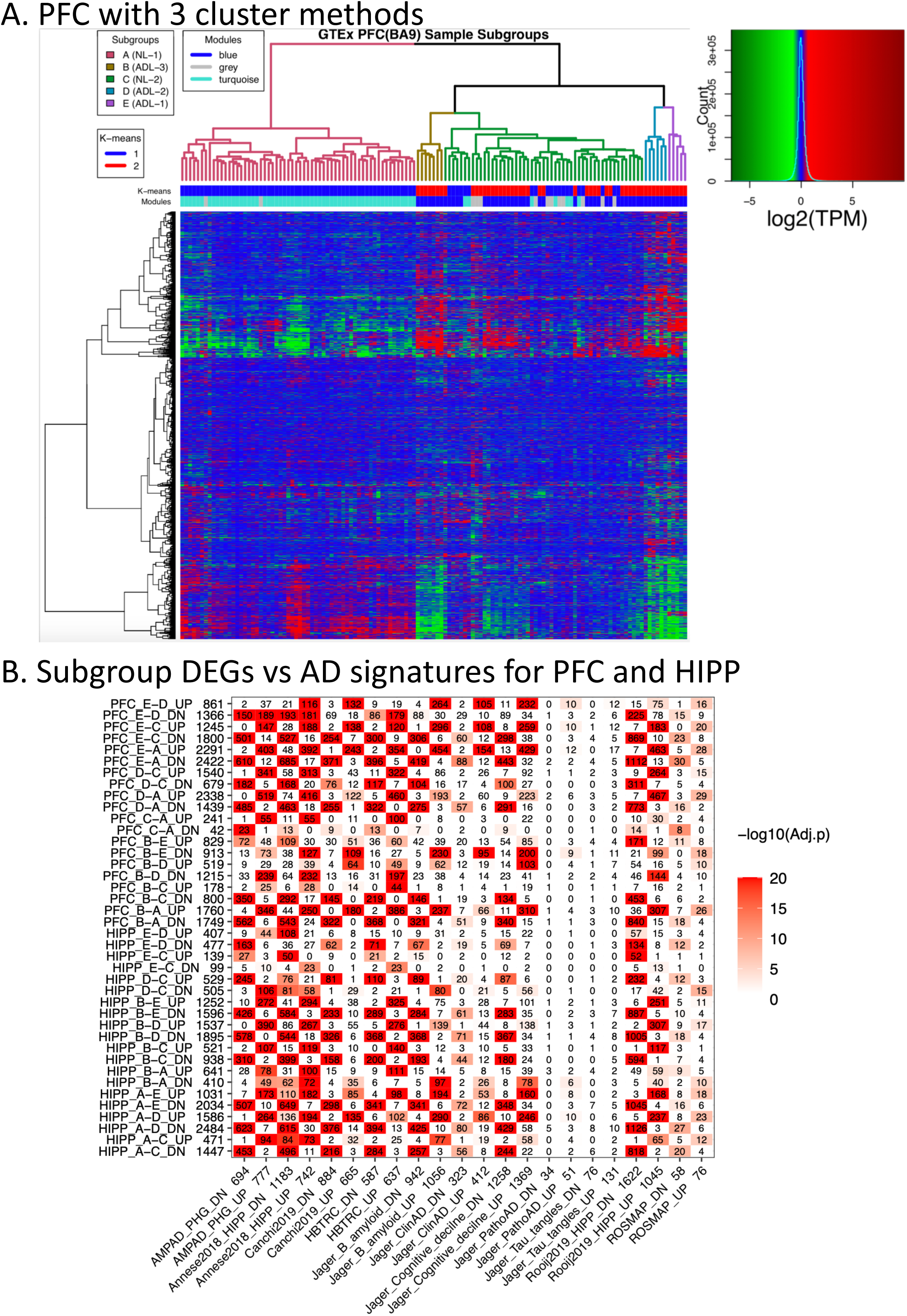
PFC with 3 cluster methods and heatmap for PFC and HIPP regions. A. 129 GTEx PFC samples show two major subtypes based on three cluster methods. B. Subgroup DEGs (fold change at 2 and FDR at 0.05) of GTEx HIPP and PFC(BA9) overlapped with reference PFC, HIPP and PHG AD signatures. Based on the direction and significance of the overlap, we were able to infer which subgroups are most AD-like and which subgroups are more healthy-like. Subgroup signatures are plotted in rows and AD signatures are plotted in columns. We separate each signature into up- and down-regulated genes and the number of genes in each signature is listed after its ID. The number in the heatmap indicates how many genes are common in the corresponding subgroup and AD signatures while the color indicates the significance of the overlap. PFC AD order: E > D, B >> C >> A. HIPP AD order: A, B >> C >> E > D.

We further performed pair-wise comparison to derive DEGs (FDR ≤ 0.05 & |fold change| > 2) between subgroups in both the PFC and HIPP regions. We compared these subgroup DEGs with our AD reference signatures (**Figure 2, Figures S3, and S4**). As shown in **Figure 2B**, DEGs derived from PFC subgroup E vs. D (denoted as PFC_E-D) significantly overlapped with most of collected AD signatures (e.g., AMPAD_PHG_DN, Rooij2019_HIPP_DN, Canchi2019_UP, Jager_B_amyloid_UP, Jager_Cognitive_decline_UP and so on) in the same direction, only 3 are reversely corelated (AMPAD_PHG_UP, HBTRC_UP and Annese2018_HIPP_UP). Furthermore, PFC_E-C, PFC_E-A showed significant overlap with AD signatures in both up- and down-regulated DEGs. PFC_B-E were reversely correlated with most of AD signatures. Based on this pattern of overlap with AD signatures, we inferred that the E subgroup was the most AD like (ADL). Since PFC_B-A, PFC_B-C, PFC_D-A, PFC_D-C all highly consistent overlapped with AD signatures in both up and down direction, we inferred that subgroups B and D were more AD-like compared to subgroups C and A. Since we did not detect reverse overlapping signatures between AD signatures and PFC_C-A DEGs, we inferred that subgroup A was the healthiest subgroup compared to subgroup C. Similarly, we considered that subgroup C was a healthier subgroup compared to B, D, and E. In summary, we inferred the order of PFC subgroups based on how similar they were to our AD signatures as E > D, B >> C > A, with E being the most AD-like and A being the most unimpaired subgroup. Since E, D, B subgroups were clustered together by all the three clustering methods (considering the top level two clusters) and they all showed high similarity with AD signatures, we considered them as the ADL subtype. For the same reason, we considered A belonged to the NL subtype. Although C was more similar to A than to B, D, E based on the order of similarity with AD signatures, due to its mixed structure, additional evidence was needed to determine if it should be classified as an NL or ADL subtype. Similarly, we inferred the order of HIPP subgroups based on how similar they were to AD signatures as A, B >> C >> E > D (**Figure 2B**). We considered A, B as the ADL subtype, E and D as the NL subtype, while C was yet to be determined.

### 3.3 The brain aging subtypes in the HIPP and PFC show a pattern of concordance that supports the spread of AD-related pathological changes from HIPP to PFC

As we have shown that both PFC and HIPP samples contained two major subtypes (NL and ADL), we then investigate if the clustering in each brain region correlates with each other. We ask that given a PFC sample being categorized as an ADL sample, will the corresponding HIPP sample (i.e., from the same donor) also cluster to a HIPP ADL subgroup, or vice versa?

To address this question, we used transcriptomics data from 98 donors whose brain tissues from both the PFC and HIPP were profiled in the GTEx. As shown in **Table 1**, **Figure 2A** and **Figure S3**, we observed that the subtype structure in the PFC and HIPP were well correlated. Particularly, we found that for the PFC ADL samples, the corresponding HIPP samples were also grouped in the ADL subtype (**Table 1**, red color samples for ADL and green color for NL samples) and none were grouped in the NL subtype. For example, For the 3 PFC_E samples, the corresponding HIPP samples were all clustered in HIPP_B, which belonged to ADL subtype. On the other hand, if samples were clustered as NL samples in HIPP, then the corresponding PFC samples were also clustered as NL samples. This pattern of correlation supports that AD-like pathological changes spread from the HIPP to the PFC as the PFC ADL subtype depended on the HIPP sample to be ADL subtype. This observation also indicated that HIPP_C should be better considered as an ADL subtype and PFC_C should be considered as an NL subtype. To make the nomenclature more meaningful, we renamed the PFC subgroups E, D, B, C and A to ADL-1, ADL-2, ADL-3, NL-2, and NL-1, respectively; so that ADL-1 was the most AD-similar while NL-1 was the most unimpaired subgroup in PFC. Similarly, we renamed the HIPP subgroups A, B, C, D, and E to ADL-1, ADL-2, ADL-3, NL-1, and NL-2 respectively.

**Table 1.** 98 overlapped samples and correlation of the subgroups of GTEx PFC and HIPP. HIPP subgroups A, B, C are ADL while E, D are NL subtype. For PFC, the subgroups E, D, B are ADL while C, A are NL subtype. As can be seen in the table, the normal brain aging subgroup: NL (green, N = 28) and AD-like subgroup: ADL (red, N = 12) are correlated across the two brain regions. HIPP AD order: A (ADL-1), B (ADL-2) >> C (ADL-3) >> E (NL-2) > D (NL-1); PFC AD order: E (ADL-1) > D (ADL-2), B (ADL-3) >> C (NL-2) > A (NL-1).

| Multi-region | PFC_E (3) | PFC_D (6) | PFC_B (3) | PFC_C (36) | PFC_A (50) |
| --- | --- | --- | --- | --- | --- |
| HIPP_A (18) | 0 | 1 | 2 | 7 | 8 |
| HIPP_B (7) | 2 | 1 | 1 | 3 | 0 |
| HIPP_C (45) | 1 | 4 | 0 | 18 | 22 |
| HIPP_E (20) | 0 | 0 | 0 | 5 | 15 |
| HIPP_D (8) | 0 | 0 | 0 | 3 | 5 |

### 3.4 DEGs between subgroups in PFC and HIPP showed significant overlap with similar function enrichment

We compared subgroup DEGs in different brain regions to evaluate if they were conserved across regions. To minimize the differences in sample size and donors between regions, we calculated DEGs from the same set of donors across regions. Specifically, we classified the controls between HIPP NL and PFC NL as the cross-region NL subtype (N = 28), overlapped HIPP ADL and PFC ADL as the cross-region ADL subtype (N = 12), and the remaining samples as the INM subtype (Intermediate samples, N = 58) (see **Table 1** for the sample counts labeled in green, red, and black, respectively). It is worth noting that the INM samples were from the ADL subgroups in HIPP, but for the corresponding PFC samples, they were classified as the PFC NL subgroups. We calculated the DEGs from pair-wise comparison of subtypes (ADL vs. NL, ADL vs. INM, and INM vs. NL) in GTEx PFC and HIPP and annotated the functions of the DEGs using DAVID tool. It is not surprising that we didn’t detect any significant DEGs in either PFC INM vs. NL or HIPP ADL vs. INM. This is because, for the INM samples in multi-region (HIPP, PFC), they are more similar to the ADL samples in HIPP but more similar to the NL samples in PFC. As shown in **Figure 3**, PFC ADL vs. NL DEGs highly overlapped with HIPP ADL vs. NL DEGs, suggesting that the gene expression changes across subgroups are largely consistent between the two brain regions. The commonly up-regulated ADL genes between PFC and HIPP are enriched for immunity, membrane, exosome, glycoprotein, and differentiation/proliferation. The commonly down-regulated ADL DEGs in PFC and HIPP are enriched for synapse/axon/dendrite, membrane, glycoprotein, neuroactive ligand-receptor interaction, ion / transport (**Table S3**). PFC specific up-regulated ADL genes are enriched for inflammatory, signal transduction, angiogenesis while HIPP specific up-regulated ADL genes are more related with cilium categories.

**Figure 3.**
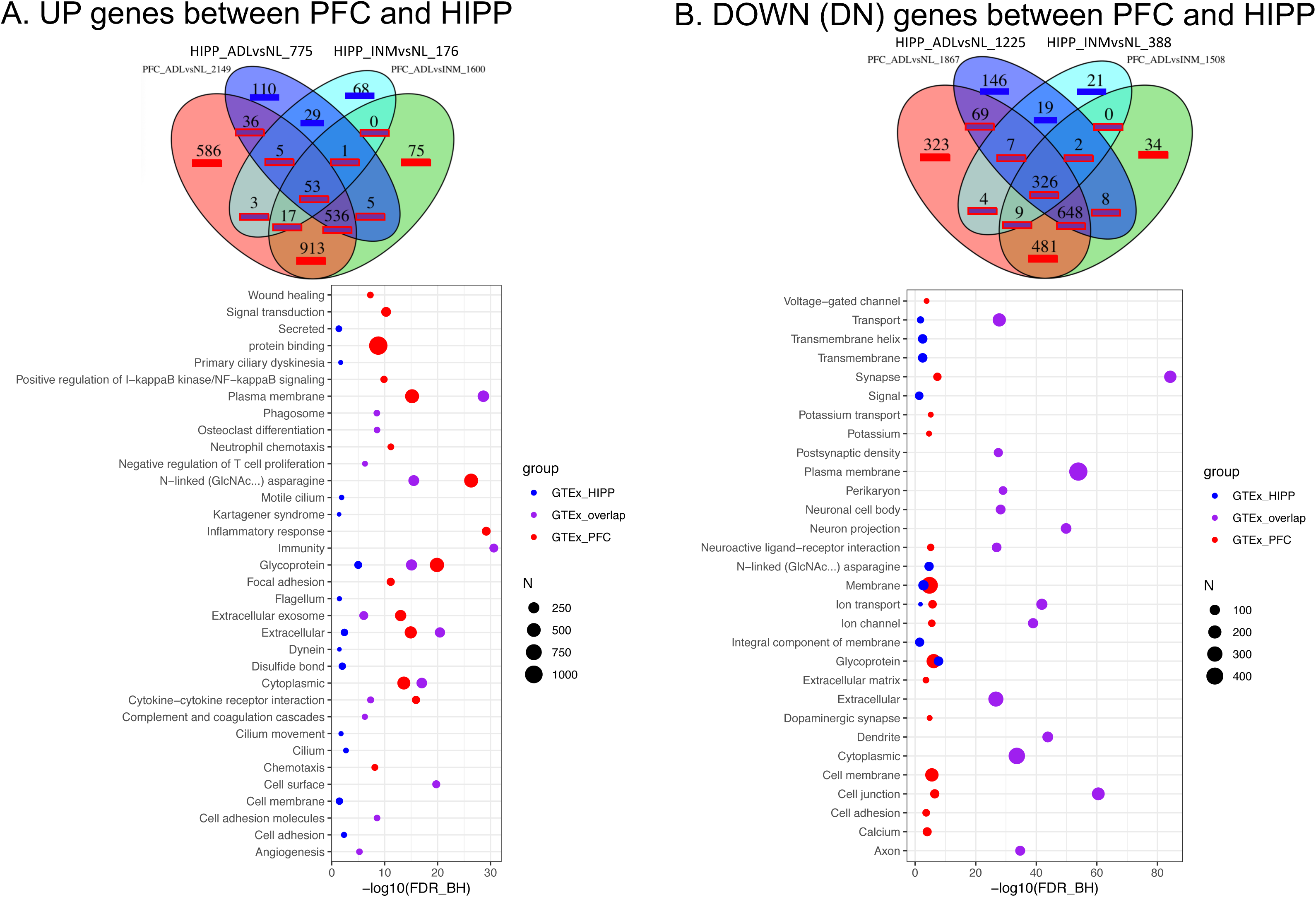
DEGs functional annotation defined in PFC and HIPP brain regions based on 3 subtypes (NL, ADL, INM). DEGs filtered with fold changes at 2 and FDR at 0.01. We list the top 15 most representative function categories with the Benjamini-Hochberg false discovery rate (FDR) < 0.05. To reduce redundancy, only one representative functional category from each identified cluster of functions was selected.

### 3.5 The brain aging subtypes are reproducible in an independent sample and can be seen in both transcriptomics and proteomics data

We recently studied the molecular features associated with brain resilience to AD; in this resilience project (RS), we profiled 167 PFC samples from cognitively unimpaired donors (**Table S1C**) for both gene and protein expression (79 MSBB and 88 ROSMAP samples) and the data has been deposited in the AD knowledge data portal [16]. Since this sample is independent from GTEx donors, we use it to evaluate if the brain aging subgroups are reproducible and if we can identify subgroups in different types of omics data.

As shown in **Figure 4**, both transcriptomics and proteomics data showed two major subtypes, which we further divided into 5 subgroups. We also calculated DEGs and DEPs between subgroups in RS transcriptomics and proteomics datasets, respectively (**Figures S5 and S6**). As shown in **Figure S5**, we rank-ordered the transcriptomic subgroups based on how similar they are to our collected AD signatures and inferred the order as C, D >> E >> A > B (C is the subgroup most similar to AD). Similarly, the rank order of the proteomic subgroups is A, C, B > E, D.

**Figure 4.**
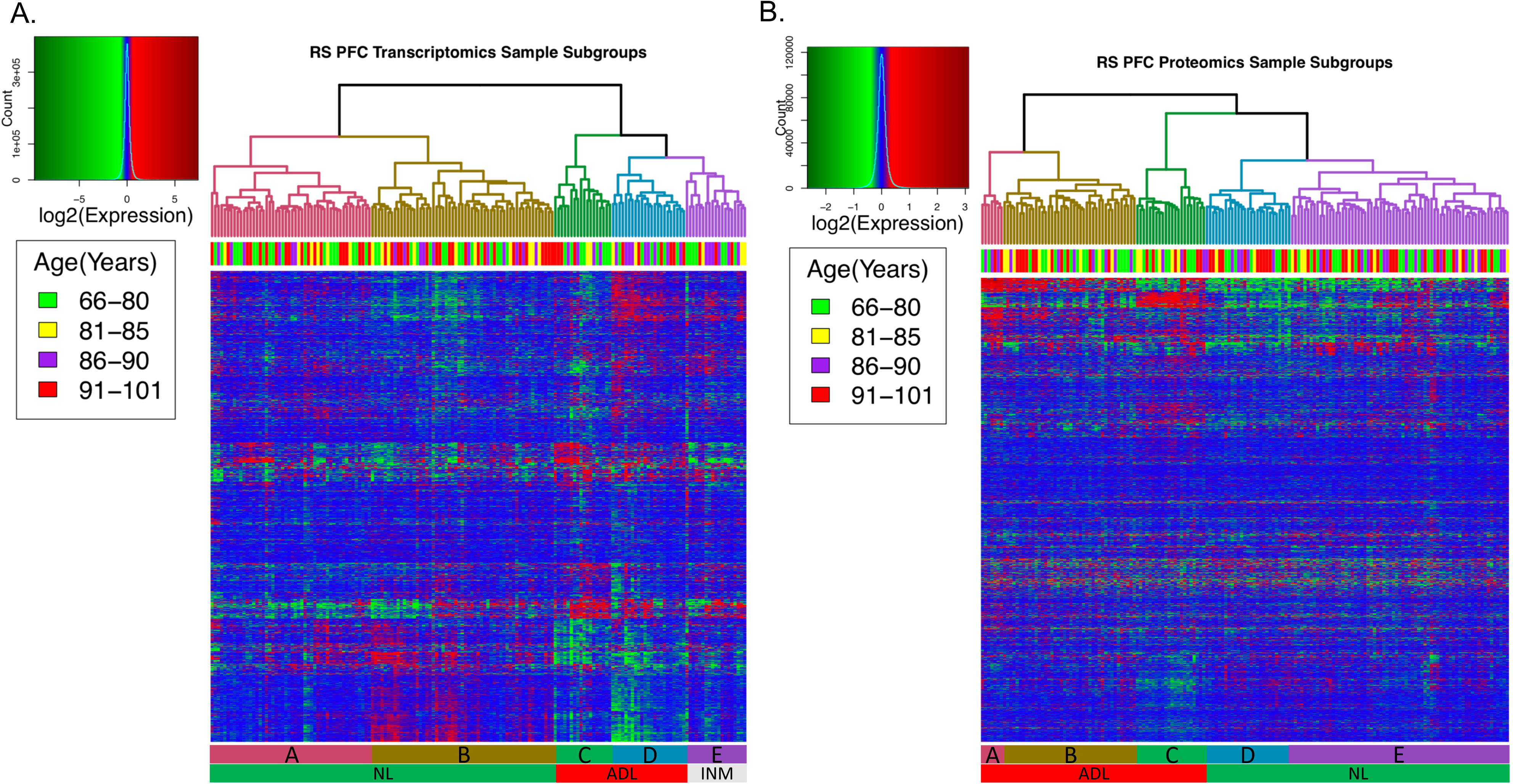
RS transcriptomics (panel A) and proteomics data (panel B) can be clustered into two major subtypes which can be further divided into 5 subgroups (A, B, C, D, E) in each type of omics data.

We also clustered samples using WSCNA and K-means and compared them with the hierarchical clusters (**Figure S6**). We observed that the WSCNA and K-means clusters were largely consistent with the hierarchical clusters. For example, most of the RS transcriptomics NL samples belonged to K-means cluster 2 and turquoise, yellow, and blue modules; while most of ADL samples belonged to K-means cluster 1 and brown and green modules. Most samples of RS proteomic subgroups A, B were contained in the K-means cluster 2 and WSCNA turquoise module (inferred as ADL subtype). RS proteomic subgroups D, E were mainly contained in K-means cluster1 and WSCNA yellow, brown, green modules (inferred as NL subtype). It is of note that we classified C as an ADL subgroup based on the order of subgroup DEGs’ similarity with our AD signatures. We compared the subtype structures defined by transcriptomics and proteomics data and observed some inconsistency between the two (**Table S4**). This raised a question as a sample might be inferred as either NL or ADL sample based on different types of omics data. To address this issue, we introduced a more stringent criterion to define NL and ADL subtypes in the RS data by creating an INM subgroup such as transcriptomic subgroup E. Unlike GTEx data (which only had NL, ADL subgroups), we classified transcriptomic subgroup E (INM-1) as an INM subgroup. As shown in **Figure S6** and **Table S4**, We defined samples overlapped between transcriptomic T_B (T NL-1), T_A (T NL-2) and proteomic P_D (P NL-1), P_E (P NL-2) as the multi-omic NL subtype, samples overlapped between transcriptomic T_C (T ADL-1), T_D (T ADL-2) and proteomic P_A (P ADL-1), P_C (P ADL-2), P_B (P ADL-3) as the ADL subtype, all other samples as the INM subtype (see **Table S4**). Based on this classification, we put samples into three subtypes, NL (normal like samples, N = 61), ADL (AD like samples, N = 21) and INM (intermediate samples, N = 85). Based on this definition, we know that RS NL and ADL samples were supported by both the transcriptomics and proteomics data.

### 3.6 GTEx vs. RS PFC subtypes DEGs showed highly similar function enrichment

We selected the RS ADL vs. NL and ADL vs. INM DEGs to compare with the corresponding DEGs in the GTEx data. As shown in **Table S5** and **Figure 5**, both GTEx and RS subtype DEGs show up-regulation of immunity, exosome and down-regulation of synapse, ion / transport genes. GTEx subtype DEGs show specific up-regulation of inflammatory and signaling pathways. RS subtype DEGs show specific up-regulation of respiratory chain, oxidative phosphorylation, neuropathy, developmental protein. This result suggests that subtype DEGs between ADL vs. NL or ADL vs. INM are largely conserved across the brain aging datasets examined in this study.

**Figure 5.**
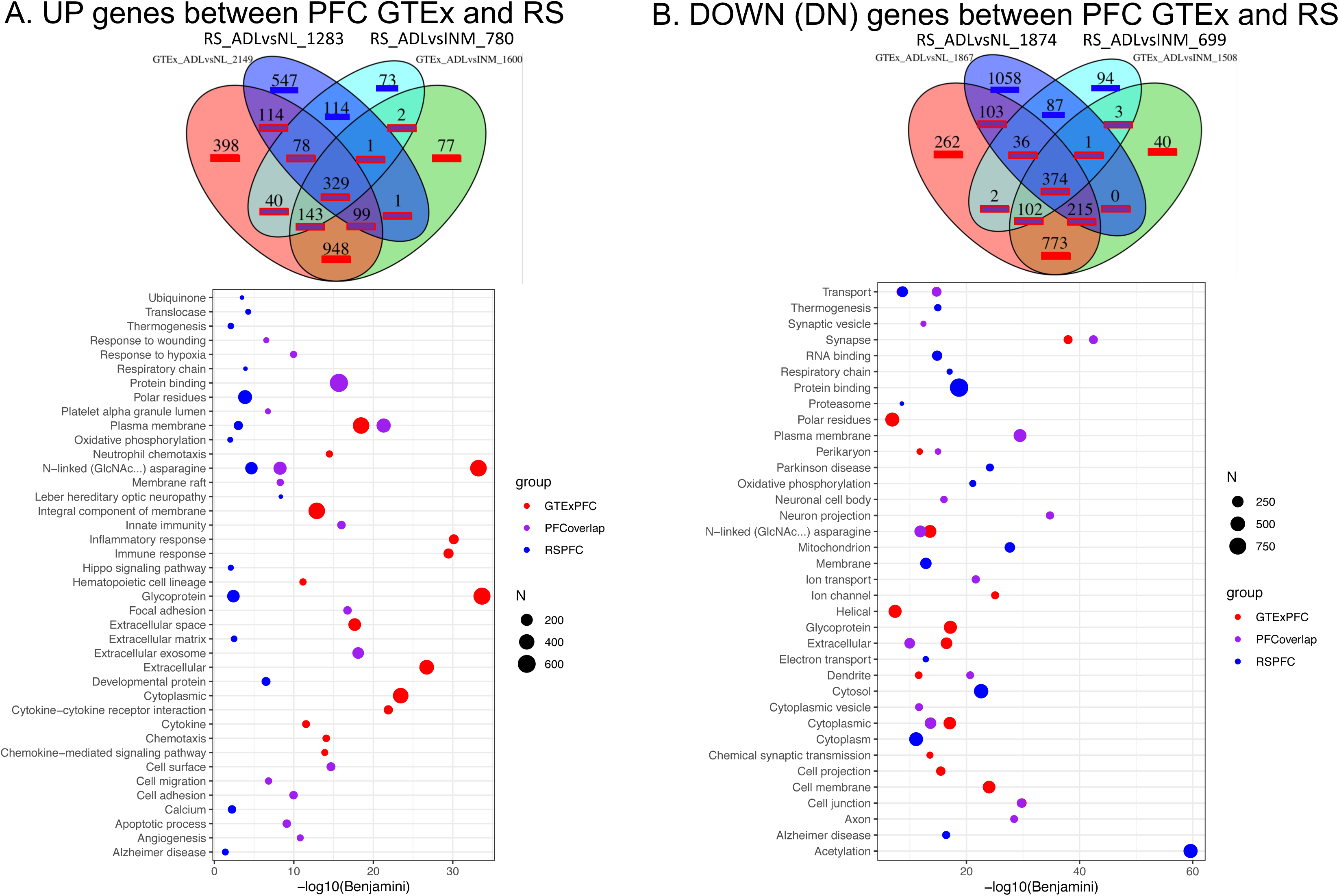
PFC RS vs. GTEx subtype comparison on function annotation with RS fold change at 1.2 and GTEx fold change at 2 and both GTEx, RS FDR ≤ 0.01. We list the most top 15 representative function categories with the Benjamini-Hochberg false discovery rate (FDR) < 0.05. To reduce redundancy, only one representative functional category from each identified cluster of functions was selected.

### 3.7 The direction of brain aging subtype DEGs and DEPs are largely consistent across brain regions and omics data types

We compared the direction of GTEx PFC DEGs with the GTEx HIPP DEGs, and the directions of RS DEGs with DEPs (all comparing ADL vs. NL subtypes). We found that for the significant DEGs in GTEx PFC and HIPP, their regulation directions are highly consistent (**Figure 6A**). For example, for DEGs at FDR ≤ 0.01, 94.8% (or 1,433 genes) showed consistent DEG directions between HIPP and PFC. It is worth noting that there are more PFC specific DEGs compared to the HIPP specific DEGs (e.g., 1,084 PFC_DN_only DEGs compared to 68 HIPP_DN_only DEGs). Similarly, the regulation directions of DEGs and DEPs are also highly consistent in the RS PFC dataset. Only a few DEGs/DEPs showed inconsistent direction (**Figure 6B** inconsistency indicated by green color). In fact, only 2 down-regulated and 26 up-regulated DEPs showed opposite regulation directions with the corresponding DEGs. The function enrichment of the consistent DEG/DEPs is very similar with the AD signatures, which includes up-regulation of host-virus interaction/innate immunity, adhesion, extracellular exosome and cytoskeleton; and down-regulation of synapse, mitochondrion, ribosome, and dendrite/axon (**Table S6**). The overall results support that the differentially expressed genes and proteins between the ADL vs. NL subtypes are largely consistent in their directions across brain regions (PFC and HIPP) and omics data types (transcriptomics and proteomics).

**Figure 6.**
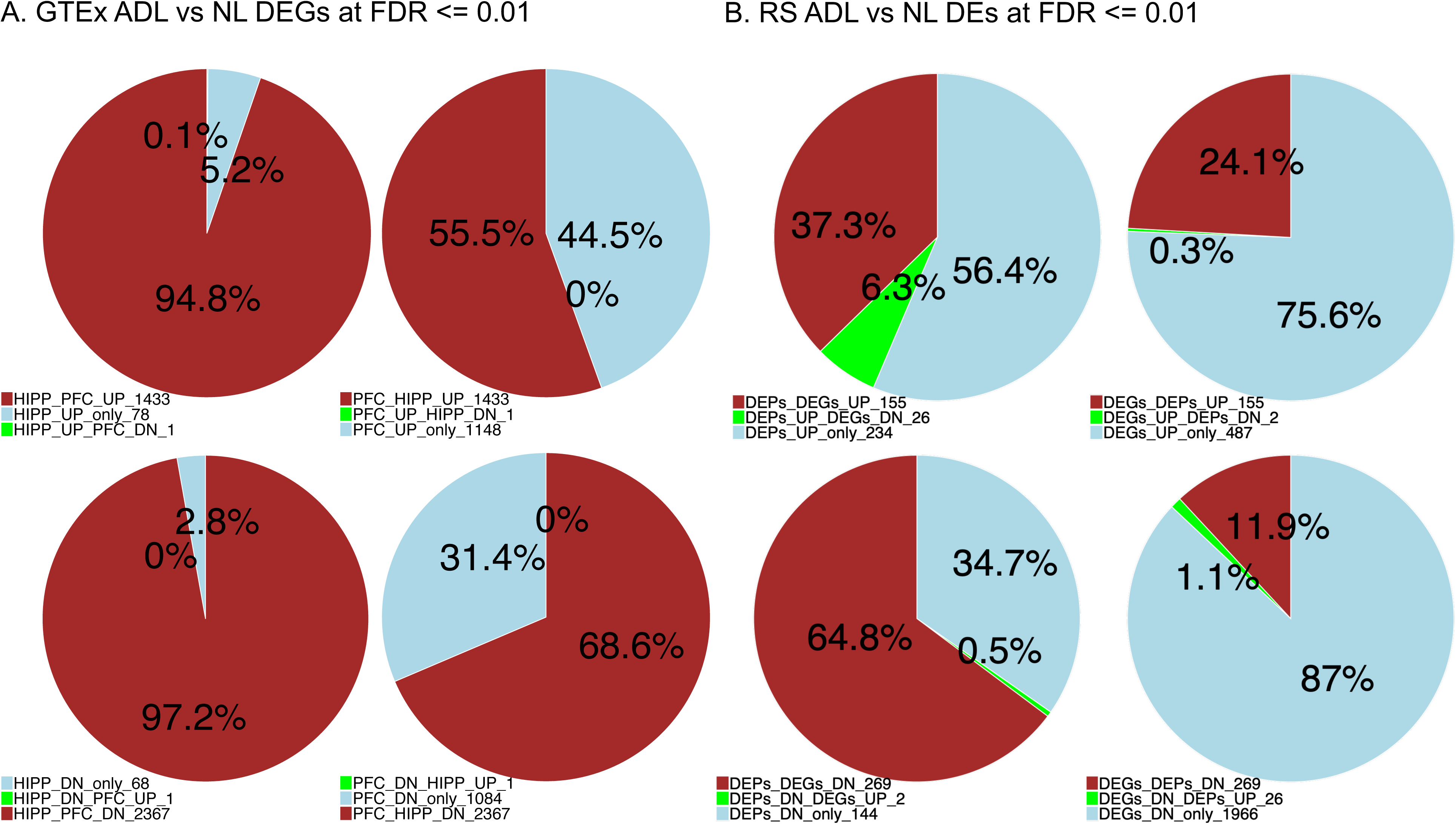
Comparison of DEGs directions across two GTEx brain regions (PFC and HIPP) and DEGs/DEPs direction in RS multi-omics data. We put the number of genes for each category at the end of the label for that group, e.g., HIPP_UP_PFC_DN_1 means there is 1 gene which is up-regulated in HIPP but down-regulated in PFC.

### 3.8 Some genes showed strong positive correlation between RS transcriptomics and proteomics

To further explore the relationship between RS transcriptomics and proteomics data, we calculated the Spearman correlation between gene and protein expression at both global and individual gene levels using transcriptomics and proteomics data from the 167 samples. We first calculated the Spearman correlation of ADL vs. NL log2FoldChanges between RNA and protein profile as shown in **Figure S7**. For all genes and proteins, spearman correlation of the gene expression fold change and the corresponding protein expression fold change between ADL vs. NL subtypes showed a positive correlation of 0.449 (p < 2.2e-16, not shown in **Figure S7**). This indicates that on the global scale, the changes of protein and mRNA levels between the two subtypes are significantly positively correlated. Interestingly, the Spearman’s correlation for the consistent differentially expressed genes (DEs_consistent: DEG and DEP’s directions are consistent) with FDR ≤ 0.01 increased to a coefficient of 0.78 (**Figure S7**). We then also calculated the Spearman correlation for each gene by directly comparing its gene expression and protein expression across samples. At FDR ≤ 0.05, we identified 235 genes that showed significant positive correlation between gene and protein expression with spearman’s correlation coefficient ρ ranging from 0.341 to 0.727. We only identified two significant negatively correlated genes, *SIRPB1* and *NDRG4* (ρ equals to -0.473 and -0.359, respectively). The function annotation showed that these genes/proteins are enriched for cytoplasm, exosome, PTMs (e.g., methylation), membrane, signal transduction, immunity, cytoskeleton, synapse (**Table S7**). The top 4 positively correlated genes are *SCIN, THNSL2, TRPV2, GSTM3*. Transient potential receptor vanilloid 2 (*TRPV2*) is a non-selective cation channel that serves as a thermo-, mechano-, and lipid sensor and have been demonstrated to have a major role in human BBB integrity[47]. Studies have shown *TRPV2* regulation during inflammation in microglia and immune cells, as well as during remyelination in oligodendrocytes. *TRPV2* has been suggested as an interesting clinical target for the development of therapeutic interventions for myelination disorders[48]. *GSTM3* colocalizes with amyloid-beta plaques in AD and reduces antioxidant defense. Additionally, *GSTM3* has been identified as a common hub of regulatory networks in blood mononuclear cells. The dysregulation of these networks may potential contribute to the development of AD[49]. *SIRPB1* has been reported as a microglial modulator of phagocytosis in Alzheimer’s disease [50].

### 3.9 The RS brain aging subtypes were more consistent across two types of omics data when significant correlated genes were used to define subtypes

We compared the subtypes derived from transcriptomics data vs. proteomics data and showed the result in **Table S4**. Interestingly, the proteomics and transcriptomics data defined subtypes appeared to have little consistency compared to the subtype-correlation derived from transcriptomics data from different brain regions as shown in **Table 1**. For example, more than half of the transcriptomic ADL-2 samples (N = 23, see **Table S4**) are categorized as proteomic NL subtypes (P_E and P_D). Using Chi-square test, the transcriptomics and proteomics subtypes are not correlated with a p-value equal to 0.474. We believe this is mainly due to the relatively low concordance between gene and protein expression data (mean of all gene Spearman’s ρ: 0.0996; min at -0.473, max at 0.727) (note this is different from the correlation between the fold change of gene and protein expression). The weak correlation between transcriptomics and proteomics has been observed in many previous studies [51, 52]. We posit that we should achieve higher concordance of the subtypes if we rely on those highly correlated proteins/genes to define the PFC brain subtypes. To test this hypothesis, we selected those DEGs or DEPs that were consistent in regulation directions and significantly correlated between gene/protein expression. At FDR ≤ 0.05, 59 genes (61 proteins) were selected with Spearman’s correlation ρ ranging between 0.341 ∼ 0.542. We used these genes/proteins’ expression to cluster the subgroups (**Figure S8**). Again, we divided samples into 5 subgroups based on their transcriptomics (T_A to T_E) and proteomics data (P_A to P_E), respectively. As shown in **Figure S9**, we inferred the rank order of transcriptomics sample subtypes for how similar they were to AD samples as B >> D > E > C > A and for proteomics subtypes as C > D > E > B, A. For this new clustering, as shown in **Table S8**, T_B, T_D and T_E were considered as T ADL-1, 2, 3 respectively, and T_A (denoted as T NL-1) was considered as an NL subgroup. For proteomics data, P_C, P_D were inferred as P ADL-1, 2 respectively, while P_A, P_B were inferred as P NL-1, 2 respectively. To get the conserved NL and ADL, we defined T_C (N = 31) and P_E (N = 31) as T INM-1, P INM-1 respectively. For the multi-omics analysis, the NL (N = 41) was defined by the samples from the common donors between T NL-1 and P NL-1, 2, ADL (N = 39) was defined by the samples from the common donors between T ADL-1, 2, 3 and P ADL-1, 2 and the rest of the samples were treated as intermediate samples (INM samples, N = 87). As shown in **Table S8**, **Figures S8, S9**, the transcriptomic and proteomic subtype structures are much more correlated compared to their correlation using all the genes. For example, we found that for the 11 P NL-1 samples that were top NL samples in proteomics (see **Table S8** green color samples), all of them were also NL samples in T NL-1, and none of them were in the 3 transcriptomics ADL subgroups. On the other hand, if samples were top ADL samples in proteomic subtypes (P ADL-1, 2, see **Table S8** red color samples), most of them were also ADL samples in transcriptomic subtypes (T ADL-1, 2, 3). A Chi-square test suggested that the ADL vs. NL subgroups defined by transcriptomics and proteomics data were significantly correlated, with a p-value < 1.32E-06.

We performed pseudo-time analysis and plot the 3D UMAP using the transcriptomics and proteomics data from two brain regions (GTEx PFC/HIPP transcriptomics and RS PFC transcriptomics and proteomics data). As shown in **Figure S10**, the NL and ADL subtype samples are well separated for the pseudo-time analysis based on GTEx PFC and RS PFC transcriptomics data. They are also separated in GTEx HIPP transcriptomics and RS PFC proteomics data, which are particularly apparent for the healthy subgroup and the most AD similar ADL subgroup. We also calculated a severity index (SI, from pseudo-time values) and **Figure S11** shows that the SIs from all four datasets were significantly different between NL and ADL subtypes.

### 3.10 Pseudo-time analysis suggests NL and ADL subtypes are different and most cognitive resilience signature genes showed differential expression between NL and ADL subtypes

We also collected and examined the expression levels of the 10 cognitive resilience (CR, with brain pathology but no dementia) genes. As shown in **Figure S12**, almost all the genes are significantly different between NL and ADL in all multi-regions and multi-comics datasets and are in the up-regulated direction in the NL subtype (except for *SH3GL1*, *ACTN4*, *UBA1* in some cases, especially for *UBA1* (down in NL)). This indicates that CR genes are associated with our brain aging subtypes.

### 3.11 GTEx Cerebellum (CRBL) also showed ADL subtype but it was different from HIPP or PFC ADL subtypes

We included the CRBL for a comparison with HIPP and PFC since it is one of the least affected brain regions by AD-related pathologies [53]; it has a specific protein expression and DNA methylation profile compared to other brain regions as well [54, 55]. The two clustering methods both suggested two major subtypes which we further divided into five subgroups (A to E) (see **Figure S13A**). The subgroup pairwise DEGs overlapped with AD reference signatures as shown in **Figure S13B** Similarly, we inferred the rank order of these subtypes for how similar they are to the AD signatures as D, C > E > A, B, and subtypes A (denoted as NL-2, N = 73 samples) and B (denoted as NL-1, N = 40 samples) were considered as NL subtype, C (denoted as ADL-2, N = 12 samples) and D (denoted as ADL-1, N = 12 samples) were considered as ADL subtype, and E (denoted INM-1, N = 28 samples) was classified as the INM subtype. Interestingly, most CRBL ADL vs. NL DEGs are up-regulated genes. For example, CRBL ADL-1 vs. NL-2 showed 960 up and 139 down-regulated genes and CRBL ADL-1 vs. NL-1 showed 1668 up and 104 down-regulated genes. The up-regulated genes significantly overlapped with up-regulated AD signatures, e.g., the DEGs between CRBL ADL-1 vs. NL-1 up (1668 genes) highly overlapped with the up-regulated AMPAD_PHG (777 genes) by 359 genes (see **Figure S13B**).

There were 80 GTEx donors with gene expression data in all the three brain regions CRBL, HIPP and PFC (see **Table S9**). Among them, 6 donors’ HIPP and PFC samples were both classified as ADL subtype in the corresponding brain region, respectively. For these 6 samples, 2 were classified as CRBL ADL (N = 12, including CRBL ADL-1, 2), 3 were classified as CRBL INM (N = 18, CRBL INM-1), and one sample was classified as CRBL NL (N = 50, including CRBL NL-1, 2). On the other hand, for CRBL ADL samples, several of them overlapped with HIPP-PFC 3 NL samples and 7 INM samples. This pattern of overlap was different from the pattern observed in the PFC ADL samples, which all overlapped with HIPP ADL samples. This suggests that the CRBL ADL subtypes could be different from the ADL subtypes in HIPP and PFC. We also conducted a cluster analysis on merged gene expression data from CRBL, PFC, HIPP samples. Our results indicate that CRBL samples are rather different from HIPP and PFC samples in their global transcriptomes (**Figure S13C**). Specifically, all CRBL samples are clustered into a separate group which is consistent with previous studies [56] (**Figure S13C**). In contrast, PFC and HIPP are mixed together to some degree, although most PFC samples are in subgroup B and most HIPP samples are in subgroups C and E (**Figure S13C**).

The function annotation of CRBL ADL vs. NL DEGs (**Table S10**) showed similar function enrichment with the HIPP/PFC ADL vs. NL DEGs (up in immunity system and down in synapse). As shown in **Table S11**, compared to HIPP, the CRBL ADL subtype has a higher number of upregulated genes compared to the number of downregulated genes. The genes that are commonly differentially expressed in both CRBL and HIPP-PFC ADL versus NL samples (fold change at 2, and FDR at 0.01) are predominantly down-regulated genes (n = 21) in synapse, plasma membrane, glycoprotein and up-regulated genes (n = 135) in plasma membrane, glycoprotein, extracellular exosome, inflammatory response and immune response. The CRBL specific up-regulated genes (n = 347) show enrichment for glycoprotein, extracellular space, inflammatory response and immune response, but there is no significant function enrichment for CRBL specific down-regulated genes.

It remains a question why CRBL contains ADL subtype while being one of the most resilient brain regions to AD neuropathology. To investigate this question, we selected a few gene sets which are representative for the biological processes tightly associated with AD and checked their gene expression levels across the three brain regions. For the up-regulated pathways we considered inflammatory response (GO:0006954) and extracellular exosome (GO:0070062); for the down-regulated pathways, we considered synapse (GO:0045202) and chemical synaptic transmission (GO:0007268). We calculated the geometric mean of gene expression of each gene set in NL and ADL samples, respectively. As shown in **Table S12**, for these four gene sets, the average percentage gene expression changes between ADL and NL subgroups in the HIPP are 13.9 %, 14%, -17.3%, -24.5%, respectively; and for the PFC they are 28.8%, 27.4%, -17.8%, and -27%, respectively. The greater percentage changes in PFC compared to HIPP is consistent with our previous observation that PFC showed a larger number of DEGs than HIPP (**Table S11**). The changes in the PFC and HIPP are much greater than the CRBL, which are 9.8%, 10.3%, -0.1%, - 1.8%, respectively. It is of note that there are almost no changes for the two down-regulated pathways (GO: synapse and GO: chemical synaptic transmission, percentage change of -0.1% and -1.8%, respectively). Furthermore, for these two down-regulated pathways, their geometric mean expression levels in the CRBL (NL at 0.932, 0.926, ADL at 0.933, 0.901, respectively) are similar to their expression levels in the PFC and HIPP ADL samples (PFC ADL: 0.902 and 0.835, respectively) and (HIPP ADL: 0.944 and 0.901, respectively), but lower than their expression levels in the PFC and HIPP NL samples (PFC NL: 1.09 and 1.107, respectively) and (HIPP NL: 1.148 and 1.234, respectively).

Taken together, these results suggest that CRBL showed ADL gene expression changes in a subset of samples. However, the changes in CRBL occurred at a much smaller scale compared to such changes in the PFC and HIPP ADL subtypes, which may explain why CRBL is more resilient to AD-related neuropathology compared to vulnerable brain regions such as the HIPP.

### 3.12 The PFC showed more DEGs between ADL vs. NL subtypes compared to HIPP mainly due to difference in gene expression in the PFC NL samples

We summarized the number of DEGs/DEPs between ADL vs. NL samples in various brain regions and datasets in **Table S13**. Interestingly, there are substantially more ADL vs. NL DEGs detected in GTEx PFC (4424 down, 4030 up) than HIPP (2042 down, 1298 up) which is consistent with the observation that PFC showed greater magnitude changes in geometric mean expression of 4 GO terms than HIPP and CRBL. It is of note that the number of samples/donors used for calculating DEGs in these two brain regions was identical for a fair comparison. The much larger number of DEGs in PFC between ADL vs. NL subtypes is to some degree unexpected since hippocampus is often considered to be more vulnerable to brain neuropathology compared to the PFC.

We are interested in finding out why there are many more DEGs observed in the PFC than in the HIPP, i.e., 1148 up-regulated and 1080 down-regulated genes specific to the PFC (FDR ≤ 0.01). For a DEG to be specific to PFC, one possible scenario is when its expression in PFC NL samples (we also call it baseline expression) is significantly different from its expression in HIPP NL samples while its expression in PFC ADL and HIPP ADL samples is similar. Using an up-regulated PFC specific DEG as an example, this may happen if this gene has a lower baseline expression level in the PFC NL samples compared to its expression in the HIPP NL samples while its expression in PFC ADL and HIPP ADL samples is similar. We call these genes as the BaseNL DEGs since the main difference is caused by the baseline gene expression (i.e., levels in the NL samples) between the two brain regions. Similarly, we define BaseADL genes if the DEG is mainly due to difference in the gene expression levels in ADL samples between the two brain regions, and BaseALL DEGs if the DEG is due to gene expression difference in both NL and ADL samples. All remaining DEGs are defined as the others type (see methods). We found that most of the PFC specific ADL vs. NL DEGs (FDR ≤ 0.01) are BaseNL genes (330 up, 789 down, see **Table S14** and **Figure S14**). Based on the enriched function of the 789 PFC BaseNL down-regulated genes, it is of note that HIPP NL samples exhibit lower expression of mitochondria, transport genes and from the function of the 330 BaseNL up-regulated genes, it suggests that HIPP NL samples have higher expression of ribosome related genes (**Table S15**). The importance of ribosome in aging has been highlighted by the research on signaling pathways and genetic screens that extend lifespan which was described in the naked mole rat, that it has a maximum-recorded lifespan of approximately 30IJyears [57].

### 3.13 Multi-region cell proportion estimated by deconvolution method suggests subtypes were associated with different proportions of major brain cell types

AD-associated gene expression changes have been shown to be highly cell-type specific and understanding the cell-type specific gene expression regulations contributing to disease development has become one of the most active topics in AD research [58–60]. We previously reported that the cellular proportions were different among brain aging subtypes in GTEx hippocampus as estimated by computational deconvolution methods [5]. Here, we applied the same methods to estimate the cellular compositions among subgroups across brain regions. As shown in **Figure 7**, the cellular compositions of neurons, oligodendrocytes, astrocytes, endothelial cells and microglia are significantly different among subgroups in both GTEx PFC and HIPP. Specifically, the PFC NL-1 and NL-2 showed higher proportions of neurons and relatively lower proportion of non-neuronal cells including the three types of glial and endothelial cells compared to the PFC ADL-1, 2, and 3. In addition, each ADL subgroup showed elevated proportion of a specific type of glial cells. For example, PFC ADL-2 mainly showed elevated proportion of microglial cells; PFC ADL-3 mainly showed increased proportion of astrocytes, while PFC ADL-1 mainly showed increased number of oligodendrocytes (**Figure 7A**). Similar to PFC, HIPP NL-2 and NL-1 showed higher proportions of neurons and lower proportion of non-neurons compared to the ADL-1, 2, 3. We also observed that each ADL subgroup had an elevated proportion of a specific type of glial cell, such as increased microglia in the HIPP ADL-2 and increased oligodendrocytes in the HIPP ADL-1 (**Figure 7B**).

**Figure 7.**
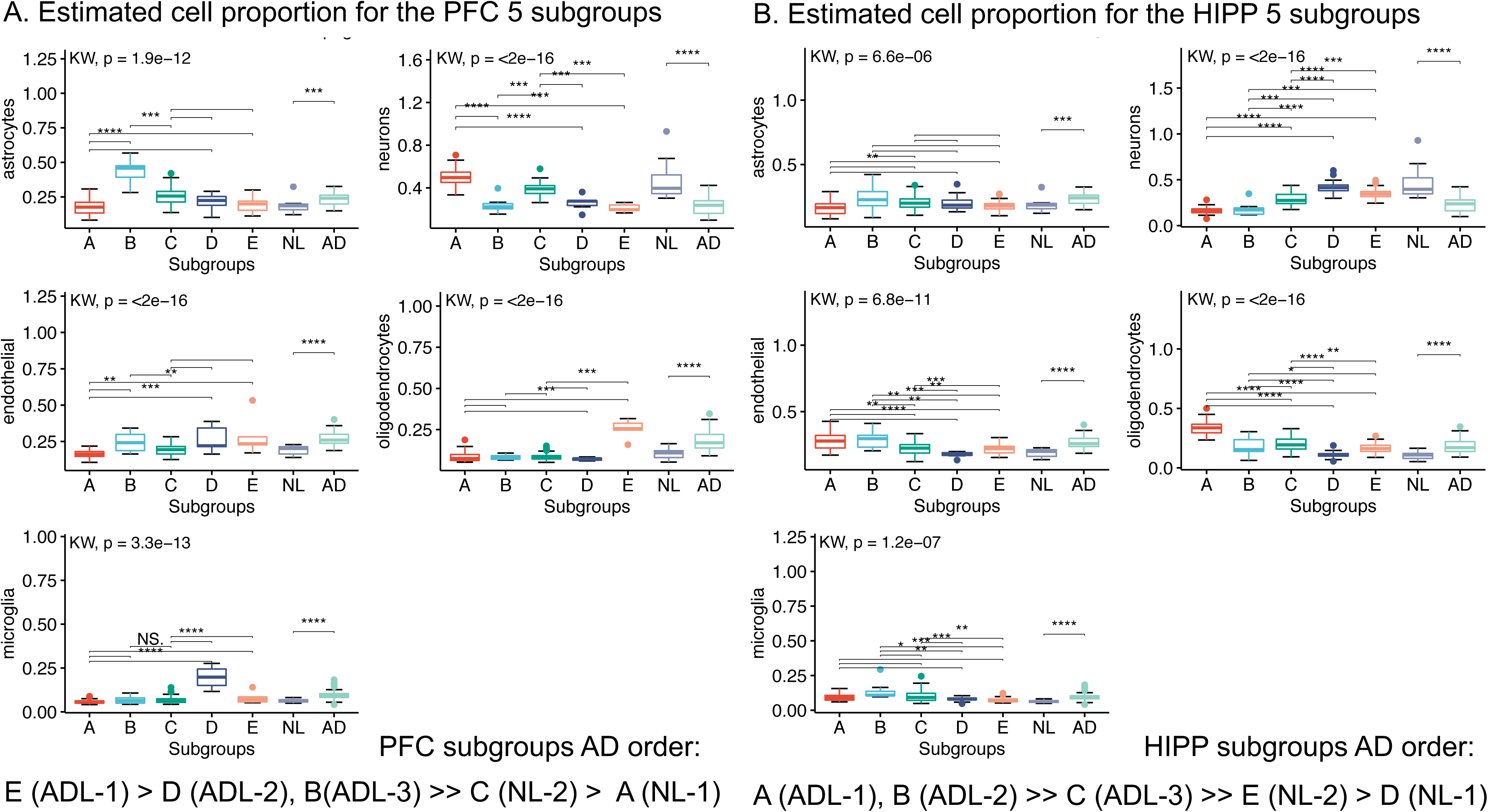
GTEx Subgroups show cell type specific proportion from DSA deconvolution analysis in each dataset. Kruskal-Wallis rank sum test (KW) and wilcox.test rank sum test were used to calculate the significance levels between the groups. Wilcox.test: "****", "***", "**", "*", "" for P value at 0, 0.0001, 0.001, 0.01, 0.05 and 1 respectively. NL: PHG (parahippocampal gyrus) NL samples, AD: PHG AD samples. DSA: Digital Sorting Algorithm.

In summary, the NL subgroups appeared to have a higher proportion of neurons and lower proportion of non-neurons (including endothelial and glial cells), suggesting cell type specificity in subgroups across brain regions. In addition, each ADL subgroup showed an elevated proportion of a specific type of glial cells in both HIPP and PFC, suggesting that the difference among ADL subgroups is at least partially driven by the cell composition difference of specific glial cell types (**Table S16**).

## 4 Discussion

Several studies have identified subtypes among aging brain samples. For instance, Peng et al. identified transcriptomics subtypes in the GTEx hippocampus (HIPP)[5], while Yang et al. conducted a multi-omics study in the prefrontal cortex (PFC)[39]. However, these studies examined a single region, limiting the scope of their findings. In this study, we compiled a large collection of multiple brain-region transcriptomics and proteomics datasets to obtain a more comprehensive understanding of the heterogeneity of brain aging and showed that certain brain subtypes could be closely related to AD. Using three clustering methods, we identified two major brain aging subtypes (NL and ADL) and these subtypes showed significant correlation across the PFC and HIPP in GTEx data. This aligns with a recent study conducted by Lee et al. [61], which analyzed multi-region transcriptomes and also reported two subtypes. In addition, the subtypes defined from transcriptomics and proteomics also showed significant correlation when a subset of well-correlated gene-protein were used to define the subgroup clustering structures. We systematically compared the brain aging subtype signatures with AD signatures and observed strong overlap with DEGs derived from comparing ADL vs. NL samples in multiple brain regions. In particular, we observed common functional enrichment such as up-regulation of exosome, inflammatory pathways and down-regulation of synapse, mitochondria for these DEGs. Different brain regions (PFC and HIPP) showed highly consistent regulatory directions, and these regulatory directions were also consistent between gene and protein expression data. Interestingly, we observed a greater number of DEGs in PFC compared to HIPP and many PFC specific DEGs were likely caused by the significant difference in baseline gene expression in NL samples between the PFC and HIPP.

It is of note that we used an age-corrected gene expression matrix as input for the hierarchical clustering. This is mainly because we did not want the clusters to simply reflect donors’ age difference which would be less interesting for the purpose of identifying the heterogeneity of brain aging. Despite age correction in data processing, we still observed some residual effect of donors’ age in subgrouping. For example, as shown in **Figure 1**, for PFC, samples from donors with age ≤ 45 were almost all clustered in NL subgroups including NL-1, 2 (PFC_A, PFC_C respectively), only 1 sample with age < 36 was clustered into ADL-3 (PFC_B); it is noteworthy that PFC_B is the weakest AD-like subgroup among all the three ADL subgroups. For HIPP, samples with age ≤ 45 were relatively evenly distributed across all NL subgroups including NL-1, 2 (HIPP_D, HIPP_E respectively) and ADL subgroups including ADL-1, 2, 3 (HIPP_A, HIPP_B, HIPP_C respectively).

We also made a comparison with the results when we did not adjust age, from which we obtained very similar clustering results (**Figure S4**). Donors with age > 45 are roughly evenly distributed across each subgroup, while all the PFC samples with age ≤ 45 being clustered in NL subgroups including NL-1, 2. For HIPP, almost all HIPP samples with age ≤ 45 were in the NL subgroups including NL-1, 2, only 1 sample with age 36-45 clustered in ADL-1 and 2 sample with age ≤45 clustered in ADL-3; it is noteworthy that ADL-3 is the weakest ADL subgroup among all 3 ADL subgroups. This result also supports that in this dataset, for some individuals (1 in ADL-1, 2 in ADL-3), the ADL gene expression changes are more likely to occur in the HIPP than the PFC.

It is worth pointing out that the clustering of the omics data is sensitive to the features used for deriving the subgroups. We showed that when we considered global protein expressions the clusters did not map well with the clusters derived from global gene expression; however, when a subset of genes/proteins were used for clustering, the subtype structures showed much stronger correlation (**Table S4 and S8**). The ideal set of features that we should use to define the subtypes is likely to be heavily problem dependent. It’s not difficult to imagine that if we try to identify subtypes that show a strong correlation with the likelihood of developing AD in the future, then a set of genes that are tightly involved in AD development would be included to the list of features to define the subtypes. On the other hand, how to best select features so the brain subtypes can optimally serve the scientific questions remains an open question and requires additional research in the future.

Our results also suggest that it is highly beneficial to consider and compare the omics data from multiple brain regions. For example, it is unexpected to see that there are more subtype DEGs in PFC than HIPP (**Table S13)**. Further investigation suggested that the difference of baseline gene expression of NL samples between PFC and HIPP could play a major role (**Table S14**). Similarly, by comparing CRBL with PFC and HIPP, we showed that for the key AD-related pathways, the gene expression changes in CRBL were at much smaller scale (**Table S12)**, supporting this brain region is among the most resilient brain regions to AD pathology. On the other hand, as all the data we have analyzed in this project were bulk tissue omics data, there is a significant need to investigate the brain aging subtypes at a single-cell level, which will help us to much better understand the cell-type specificity that drive the brain aging subtype, such a cellular level of understanding is truly needed for us to elucidate how brain-aging and AD are connected.

In summary, our research indicates that there are distinct molecular changes in the brain that are associated with healthy aging (NL samples) and Alzheimer’s disease-like aging (ADL samples). Specifically, we found that the NL subtype was characterized by low levels of non-neurons (including endothelial and glial cells) and high levels of neurons, with increased mitochondrial, synaptic abundance in neurons and decreased inflammation in glial cells. In contrast, ADL samples exhibited the opposite patterns of these changes. Our findings highlight the importance of studying the post-transcriptional regulation of proteins in aging, as the transcriptome alone does not capture the full spectrum of changes in the human aging brain. In addition, the investigation of normative aging with the molecular subtype ADL offers the possibility of identifying the earliest molecular changes (e.g., ribosome changes) associated with preclinical AD, which may lead to the identification of novel biomarker candidates (e.g., from cerebrospinal fluid or blood) and therapeutic targets in the future.

## Consent Statement

All ROSMAP participants enrolled without known dementia and agreed to detailed clinical evaluation and brain donation at death. Both studies were approved by an Institutional Review Board of Rush University Medical Center. Each participant signed an informed consent, Anatomic Gift Act, and repository consent to allow their data to be repurposed.

## Data Availability Statement

Publicly datasets were analyzed in this study: Genotype-Tissue Expression (GTEx) data (https://www.gtexportal.org/home/datasets), and the human postmortem sequencing data in AD Knowledge Portal (https://adknowledgeportal.synapse.org) which include: (1) Resilience project data (Synapse ID: syn52676458) including the Mount Sinai Brain Bank (MSBB) and the Religious Orders Study and Memory and Aging Project (ROSMAP) and (2) parahippocampal gyrus dataset (PHG, Synapse ID: syn20818651).

## Author Contributions

ZT conceived the main concept of the work. ZT and SP contributed to conceptualization, methodology, analysis, and preparation of the manuscript. SP, EW, MW, XW, SP, DMH, JP, and BZ helped to discuss and improve the work. All authors have read and approved the manuscript for submission.

## Supplementary

### Supplementary Figures

**Figure S1.** Proteomics gene lists in Frontal Cortex. **A.** Proteomics: Cognitive trajectory stability (columns) vs. AD (rows) Signatures in Frontal Cortex. AD signatures are plotted in rows and cognitive trajectory stability signatures are plotted in columns. **B.** Reanalyzed the Ping2020 data with 5 hierarchical clusters and 4 WSCNA modules. **C.** Reanalyzed the Ping2020 data for 5 hierarchical clusters (without AD samples) and 4 WSCNA modules. **D.** Selected AD Transcriptomic (columns) vs. Proteomic (rows) Signatures in Front Cortex. AD proteomic signatures are plotted in rows and transcriptomic signatures are plotted in columns. We separate each signature into up- and down-regulated genes and the number of genes in each signature is listed after its ID. The number in the heatmap indicates how many genes are common in the corresponding AD proteomic and AD transcriptomic signatures while the color indicates the significance of the overlap. Ping2020_BA9_AsymNL AD signatures is from the differentially expressed proteins between Diagnosis AsymAD samples vs. Diagnosis control samples. We separate each signature into up- and down-regulated genes and the number of genes in each signature is listed after its ID. The number in the heatmap indicates how many genes are common in the corresponding cognitive trajectory stability and AD signatures while the color indicates the significance of the overlap.

**Figure S2.** The Venn plot and function annotation of global transcriptomic and proteomic AD signatures (TP_AD). We list the top 15 most representative function categories with the Benjamini-Hochberg false discovery rate (FDR) < 0.05. To reduce redundancy, only one representative functional category from each identified cluster of functions was selected.

**Figure S3.** 129 GTEx HIPP samples show two major subtypes based on three cluster methods.

**Figure S4.** Multi-region (HIPP and PFC) with no Age adjusted shows two major subtypes based on 5 subgroups.

**Figure S5.** Subgroups DEs (FDR at 0.01) from 167 overlapped RS samples between transcriptomics and proteomics overlapped with reference PFC AD signatures. AD order in DEGs: C, D > E >> A > B. AD order in DEPs: A, C, B > E, D. T_AD for transcriptomic AD signatures, P_AD for proteomic AD signatures.

**Figure S6.** Multi-Omics (RS gene and protein) shows two major subtypes based on three cluster methods. **A.** RS transcriptomic subgroup D and C are consistent with most K-means cluster 1 and most WSCNA module green and module brown samples (we inferred as ADL subtype) but there are also few samples in K-means cluster 1 and WSCNA brown and green modules in RS transcriptomics subgroup A and B (we inferred as NL subtype). **B.** RS proteomic subgroups A, B and C are mainly contained in the alternative method K-means cluster 2 and WSCNA module turquoise (we inferred as ADL subtype). RS proteomic subgroup D, E are also mainly contained in K-means cluster1 and the WSCNA module yellow, brown, green (we inferred as NL subtype).

**Figure S7.** The Spearman’s correlation of ADL vs. NL log2FoldChanges between RNA and protein profile for genes among different DE categories (FDR ≤ 0.01). DEs_consistent: DEPs and DEGs directions are consistent, DEs_reverse: DEPs and DEGs have opposite directions. DEGs_only: genes with FDR ≤ 0.01 but proteins with FDR > 0.01. DEPs_only: proteins with FDR ≤ 0.01 but genes with FDR > 0.01. DEs_0.05: 0.01 < FDR of DEGs and DEPs ≤ 0.05. UN_DEs: the genes with FDR > 0.05 in both transcriptomics and proteomics. R correlation for each category is listed in the upper left corner.

**Figure S8.** Multi-Omics (RS gene and protein) 167 sample based consistent DEs and significant correlative genes (Num. genes = 59) shows two major subtypes based on 5 subgroups. **A.** Subtype NL (including subgroup A), Subtype ADL (including subgroup B, E, D); **B.** Subtype NL (including B and A), Subtype ADL (including subgroup C, D).

**Figure S9.** RS 167 sample based consistent DEs and significant correlative genes (Num. genes = 59) pair-wise subgroup DEs overlapped with AD signatures. AD order in DEGs: B >> D > E > C > A. order in DEPs: C > D > E > B, A. T_AD for transcriptomic AD signatures, P_AD for proteomic AD signatures.

**Figure S10.** GTEx and RS Subtypes in 3D UMAP plot.

**Figure S11.** GTEx and RS datasets 3 Subtypes in pseudotime plot. Wilcox.test rank sum test were used to calculate the significance levels between AD_Aging and Healthy_Aging. Wilcox.test: "****", "***", "**", "*", "" for P at 0, 0.0001, 0.001, 0.01, 0.05 and 1 respectively.

**Figure S12.** 10 CR genes/proteins expression boxplot in GTEx and RS 4 datasets. **A.** 10 CR genes in HIPP GTEx transcriptomics. **B.** 10 CR genes in GTEx PFC transcriptomics. **C.** 10 CR genes in PFC RS proteomics. **D.** 10 CR genes in PFC RS transcriptomics. N for NL, I for INM, A for ADL. Kruskal-Wallis rank sum test (KW) and wilcox.test rank sum test were used to calculate the significance levels between the groups. Wilcox.test: "****", "***", "**", "*", "" for P at 0, 0.0001, 0.001, 0.01, 0.05 and 1 respectively. KW test: P value in top left of each sub-figure.

**Figure S13.** GTEx CRBL subgroups and subgroups DEGs vs AD signatures. **A.** GTEx CRBL shows two major subtypes based on 5 subgroups. Aging_SubType: GTEx CRBL NL, ADL, INM subtype. **B.** CRBL pair-wised subgroup DEGs (|Fold change| > 2, FDR < 0.05) overlapped with AD signatures. AD order in DEGs: D, C > E > A, B. **C.** GTEx CRBL, PFC, HIPP subgroups.

**Figure S14.** Top PFC specific ADL vs. NL DEGs (FDR ≤ 0.01) average expression after adjusted covariances. A is for ADL subtype, I is for INM subtype, N is for NL subtype, F is for PFC region, H is for HIPP region. E.g., A_F is for ADL subtype in PFC region. UP.NL or DN.NL is for BaseNL genes due to NL samples gene expression baseline levels. UP.ADL or DN.ADL is for up or down-regulated BaseADL genes due to ADL samples gene expression baseline levels. UP.ALL is for up-regulated gene due to both NL and ADL samples gene expression baseline levels. Kruskal-Wallis rank sum test (KW) and wilcox.test rank sum test were used to calculate the significance levels between the groups. Wilcox.test: "****", "***", "**", "*", "" for P at 0, 0.0001, 0.001, 0.01, 0.05 and 1 respectively.

### Supplementary Tables

**Table S1.** Description of datasets and aging and AD gene signatures and function annotation of global signatures.

**Table S2.** The best number of subgroups for each dataset by 2 validation criteria type.

**Table S3.** The function annotation of GTEx PFC and HIPP 3 pair-wise subtype DEGs (ADL vs. NL, ADL vs. INM, INM vs. NL).

**Table S4.** 167 overlapped RS samples in PFC region with consistent NL (green, N = 61) and ADL (red, N = 21) subtype. DEPs AD order: A (ADL-1), C (ADL-2), B (ADL-3) >> E (NL-2), D (NL-1); DEGs AD order: C (ADL-1), D (ADL-2) > E (INM-1) >> A (NL-2) > B (NL-1).

**Table S5.** The function annotation of GTEx PFC and RS 3 pair-wise subtype DEGs (ADL vs. NL, ADL vs. INM, INM vs. NL).

**Table S6.** The function annotation of consistent direction genes in GTEx and RS data.

**Table S7.** The function annotation of 235 RS significant correlated genes.

**Table S8.** Comparison of 167 overlapped RS samples in transcriptomics data derived subtypes and proteomics data derived subtypes based on consistent and significant mRNA-protein correlated genes (Num. genes = 59): The ADL subtypes (N = 39) are highlighted in red color and NL subtypes (N = 41) are highlighted in green color. RS transcriptomics AD order: B (ADL-1) >> D (ADL-2) > E (ADL-3) > C (INM-1) > A (NL-1) and RS proteomics AD order as C (ADL-1) > D (ADL-2) > E (INM-1) > B (NL-2), A (NL-1).

**Table S9.** 80 CRBL subgroups samples overlapped with HIPP-PFC NL and ADL samples.

**Table S10.** The function annotation of CRBL ADL vs. NL DEGs.

**Table S11.** A summary list of numbers of DEGs between 3 subtypes derived from PFC-HIPP multi-region and CRBL data.

**Table S12.** Geometric mean expression of four representative pathway genes in NL and ADL samples across CRBL, PFC, and HIPP.

**Table S13.** A summary list of numbers of DEGs or DEPs between subgroups derived from multi-region and multi-omics data.

**Table S14.** PFC specific ADL vs. NL DEGs (FDR ≤ 0.01) can be divided into several group including BaseNL (different NL gene expression, no significant difference in ADL samples between PFC and HIPP), BaseADL (different ADL gene expression but no significant difference in NL samples between PFC and HIPP), BaseALL (different gene expression in both NL and ADL samples), and Others (all other situations). The number of significant DEGs are listed in the table.

**Table S15.** The function annotation of 330 up-regulated and 789 down-regulated BaseNL genes.

**Table S16.** Glial and Endothelial cell type proportion in AD-like subgroup.

