## Supplemental Figures and Tables for "Human brain aging heterogeneity observed from multi-region omics data reveals a subtype closely related to Alzheimer’s disease"

**Figure S1.** Proteomics gene lists in Frontal Cortex. **A.** Proteomics: Cognitive trajectory stability (columns) vs. AD (rows) Signatures in Frontal Cortex. AD signatures are plotted in rows and cognitive trajectory stability signatures are plotted in columns. **B.** Reanalyzed the Ping2020 data with 5 hierarchical clusters and 4 WSCNA modules. **C.** Reanalyzed the Ping2020 data for 5 hierarchical clusters (without AD samples) and 4 WSCNA modules. **D.** Selected AD Transcriptomic (columns) vs. Proteomic (rows) Signatures in Front Cortex. AD proteomic signatures are plotted in rows and transcriptomic signatures are plotted in columns. We separate each signature into up- and down- regulated genes and the number of genes in each signature is listed after its ID. The number in the heatmap indicates how many genes are common in the corresponding AD proteomic and AD transcriptomic signatures while the color indicates the significance of the overlap. Ping2020_BA9_AsymNL AD signatures is from the differentially expressed proteins between Diagnosis AsymAD samples vs. Diagnosis control samples. We separate each signature into up- and down- regulated genes and the number of genes in each signature is listed after its ID. The number in the heatmap indicates how many genes are common in the corresponding cognitive trajectory stability and AD signatures while the color indicates the significance of the overlap.

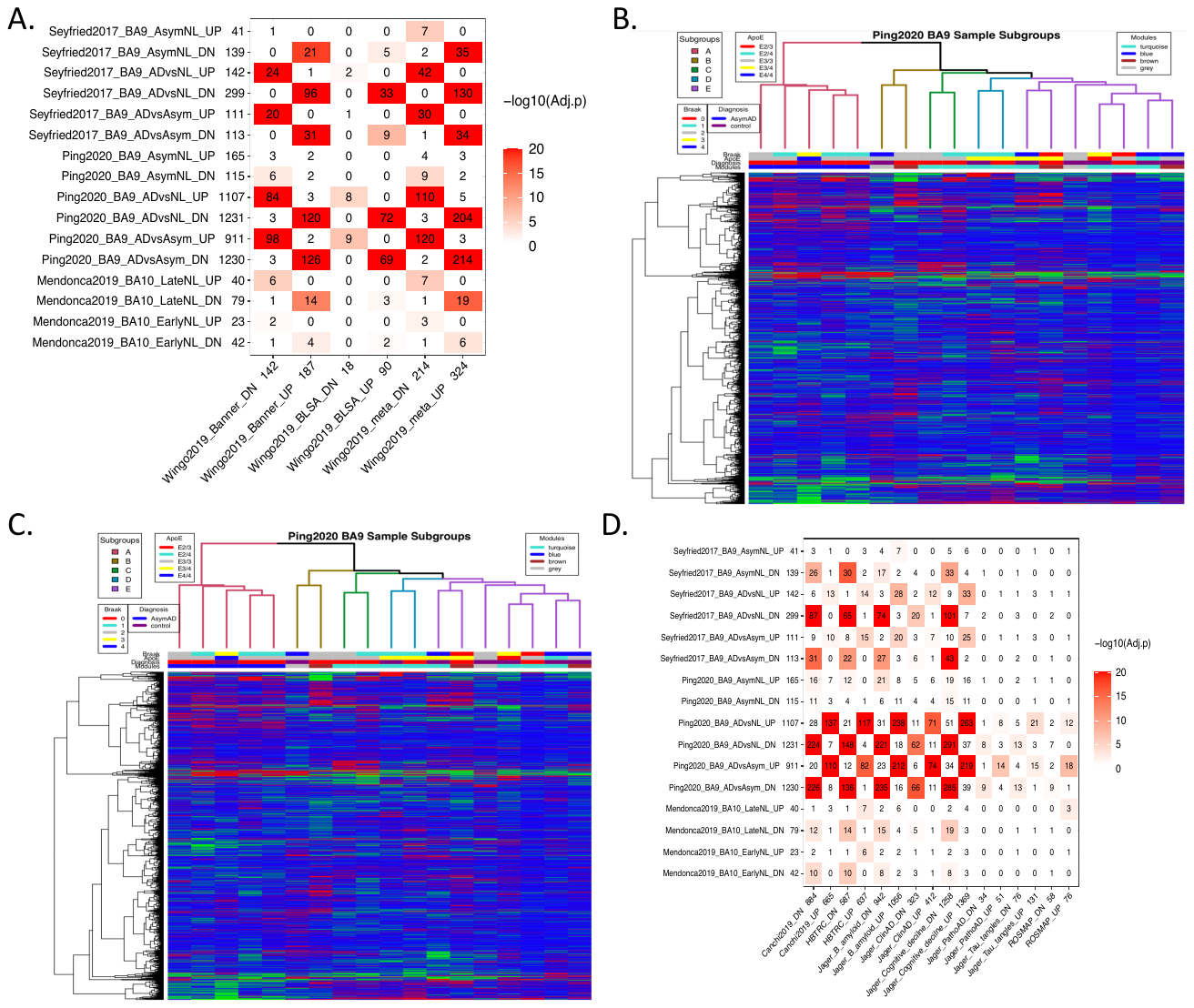

**Figure S2.** The Venn plot and function annotation of global transcriptomic and proteomic AD signatures (TP_AD). We list the top 15 most representative function categories with the Benjamini-Hochberg false discovery rate (FDR) < 0.05. To reduce redundancy, only one representative functional category from each identified cluster of functions was selected.

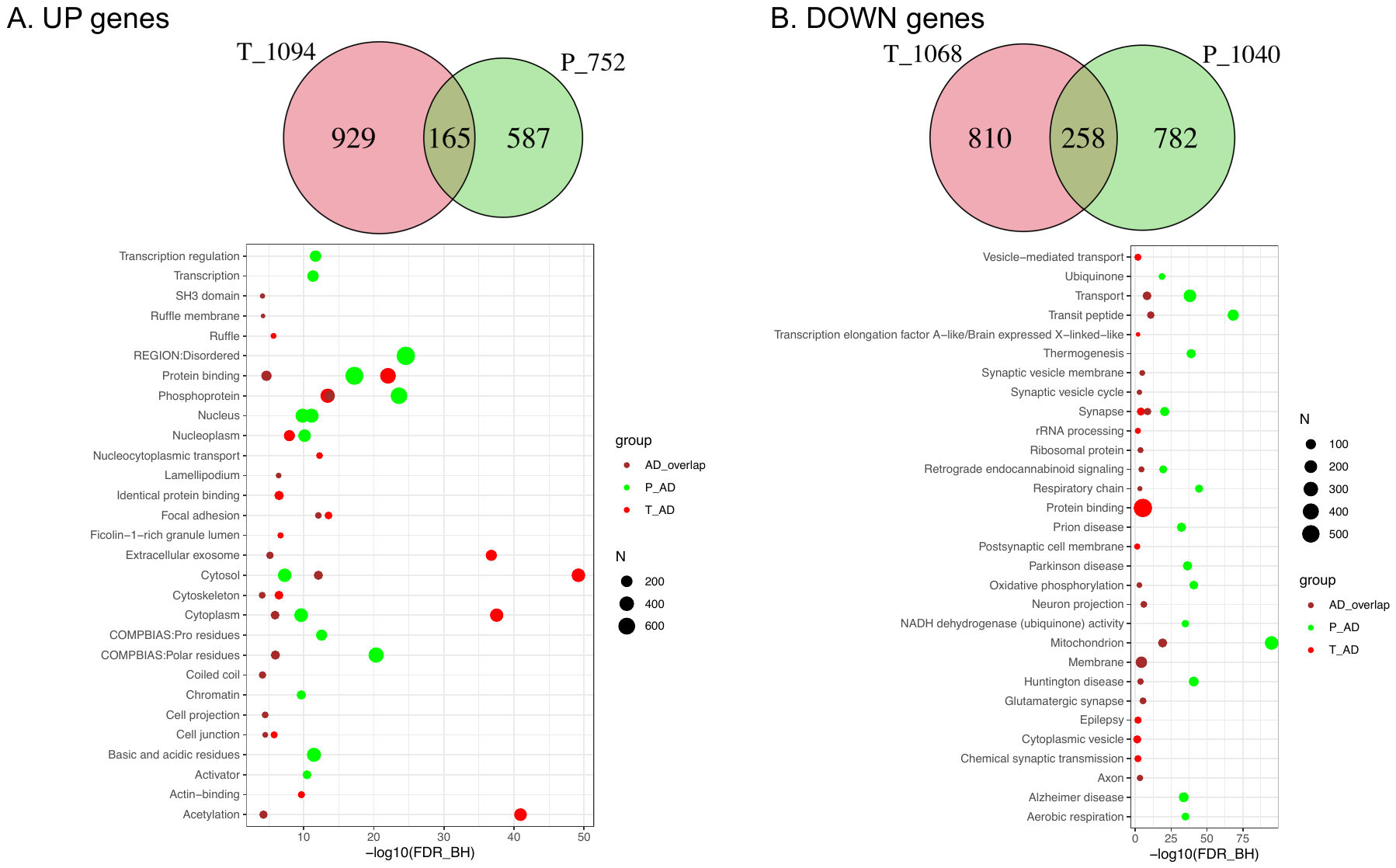

**Figure S3.** 129 GTEx HIPP samples show two major subtypes based on three cluster methods.

**Figure S4.** Multi-region (HIPP and PFC) with no Age adjusted shows two major subtypes based on 5 subgroups.

**Figure S5.** Subgroups DEs (FDR at 0.01) from 167 overlapped RS samples between transcriptomics and proteomics overlapped with reference PFC AD signatures. AD order in DEGs: C, D > E >> A > B. AD order in DEPs: A, C, B > E, D. T_AD for transcriptomic AD signatures, P_AD for proteomic AD signatures.

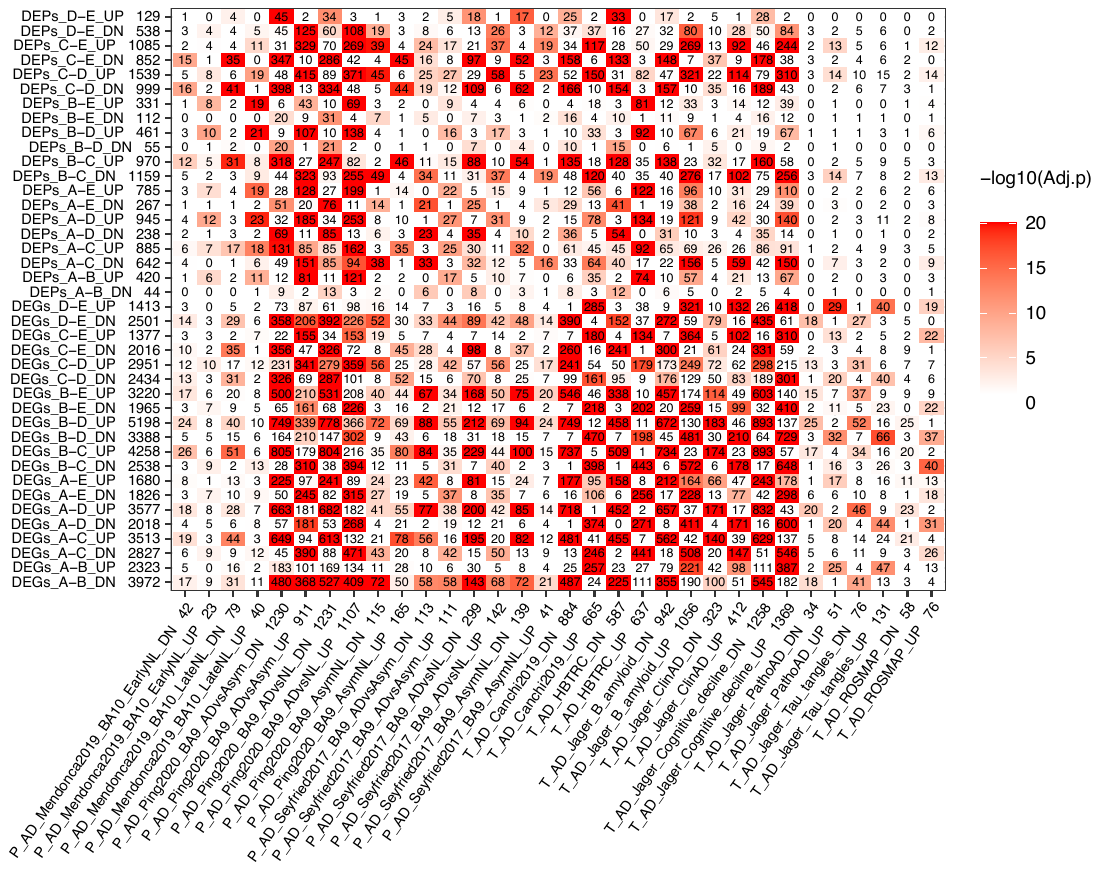

**Figure S6.** Multi-Omics (RS gene and protein) shows two major subtypes based on three cluster methods. **A.** RS transcriptomic subgroup D and C are consistent with most K-means cluster 1 and most WSCNA module green and module brown samples (we inferred as ADL subtype) but there are also few samples in K-means cluster 1 and WSCNA brown and green modules in RS transcriptomics subgroup A and B (we inferred as NL subtype). **B.** RS proteomic subgroups A, B and C are mainly contained in the alternative method K-means cluster 2 and WSCNA module turquoise (we inferred as ADL subtype). RS proteomic subgroup D, E are also mainly contained in K-means cluster1 and the WSCNA module yellow, brown, green (we inferred as NL subtype).

**Figure S7.** The Spearman’s correlation of ADL vs. NL log2FoldChanges between RNA and protein profile for genes among different DE categories (FDR ≤ 0.01). DEs_consistent: DEPs and DEGs directions are consistent, DEs_reverse: DEPs and DEGs have opposite directions. DEGs_only: genes with FDR ≤ 0.01 but proteins with FDR > 0.01. DEPs_only: proteins with FDR ≤ 0.01 but genes with FDR > 0.01. DEs_0.05: 0.01 < FDR of DEGs and DEPs ≤ 0.05. UN_DEs: the genes with FDR > 0.05 in both transcriptomics and proteomics. R correlation for each category is listed in the upper left corner.

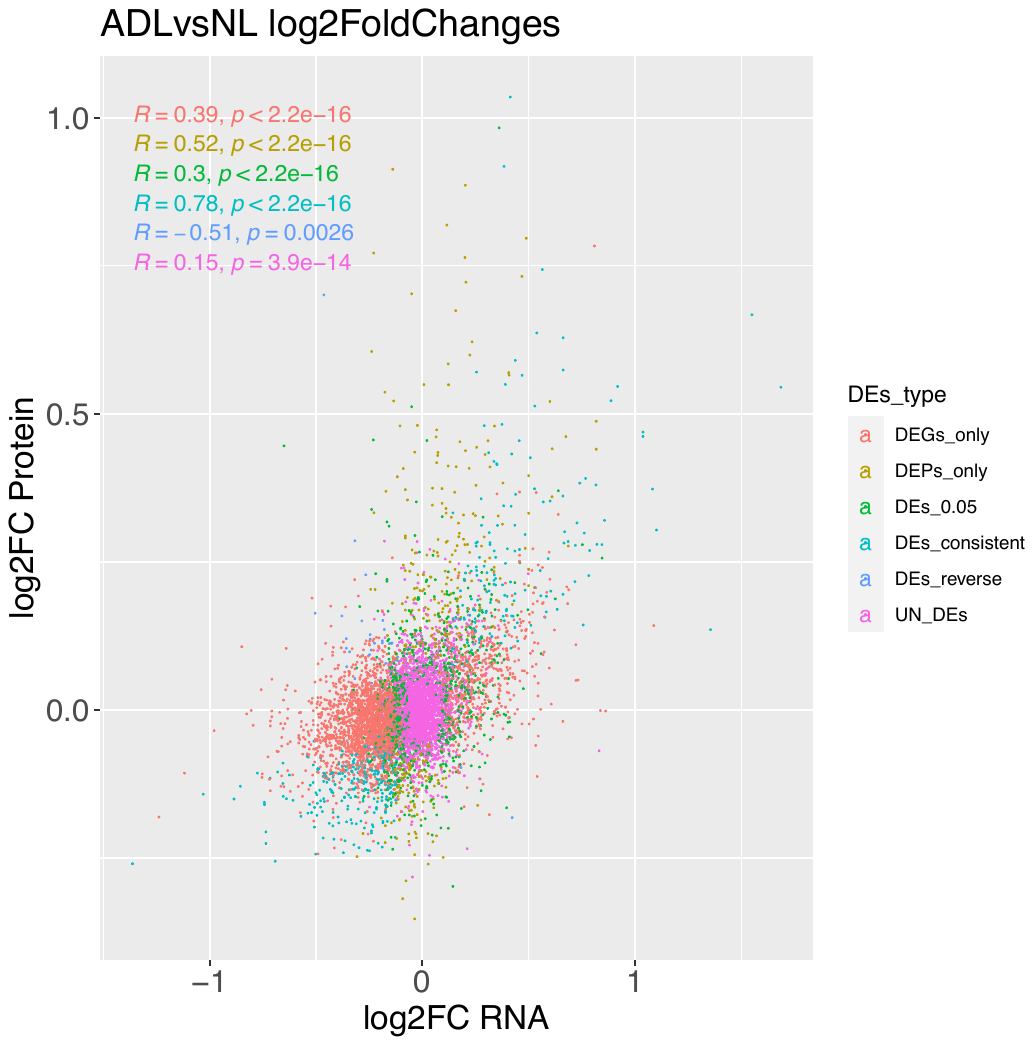

**Figure S8.** Multi-Omics (RS gene and protein) 167 sample based consistent DEs and significant correlative genes (Num. genes = 59) shows two major subtypes based on 5 subgroups. **A.** Subtype NL (including subgroup A), Subtype ADL (including subgroup B, E, D); **B.** Subtype NL (including B and A), Subtype ADL (including subgroup C, D).

**Figure S9.** RS 167 sample based consistent DEs and significant correlative genes (Num. genes = 59) pair-wise subgroup DEs overlapped with AD signatures. AD order in DEGs: B >> D > E > C > A. order in DEPs: C > D > E > B, A. T_AD for transcriptomic AD signatures, P_AD for proteomic AD signatures.

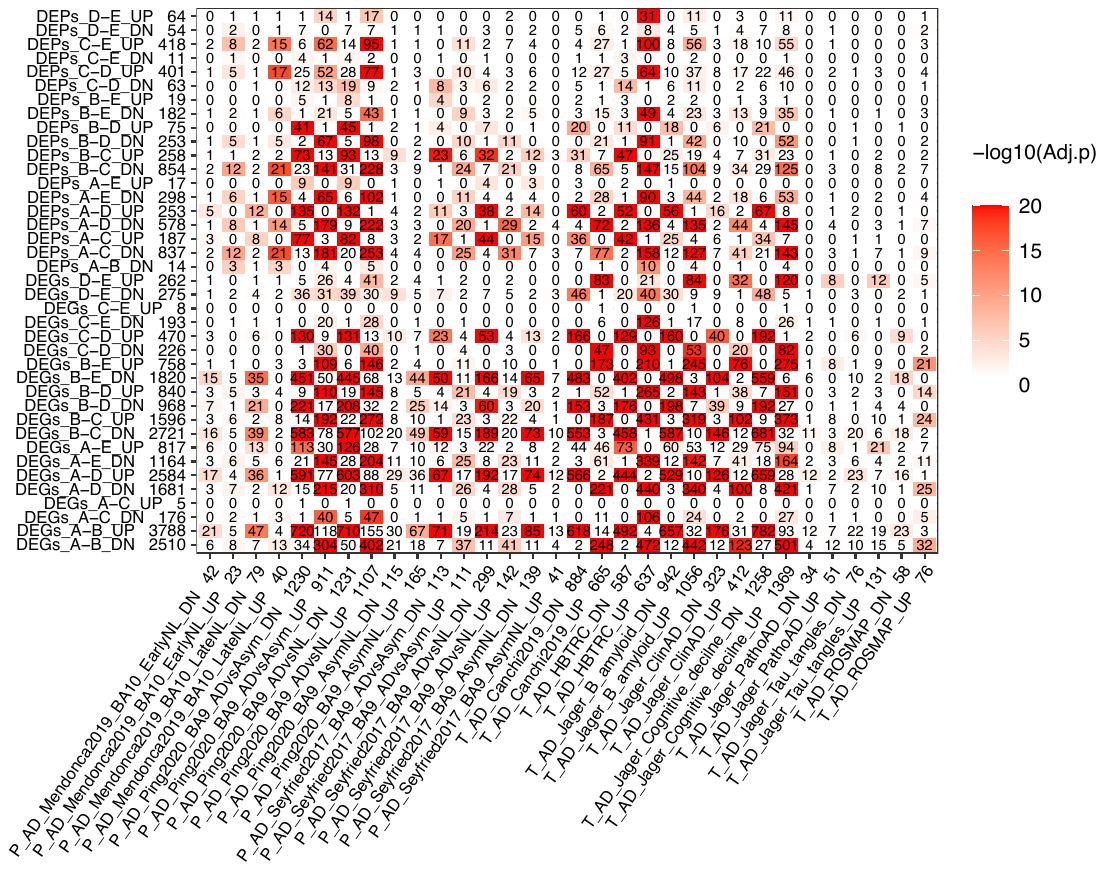

**Figure S10.** GTEx and RS Subtypes in 3D UMAP plot.

**Figure S11.** GTEx and RS datasets 3 Subtypes in pseudotime plot. Wilcox.test rank sum test were used to calculate the significance levels between AD_Aging and Healthy_Aging. Wilcox.test: "****", "***", "**", "*", "" for P at 0, 0.0001, 0.001, 0.01, 0.05 and 1 respectively.

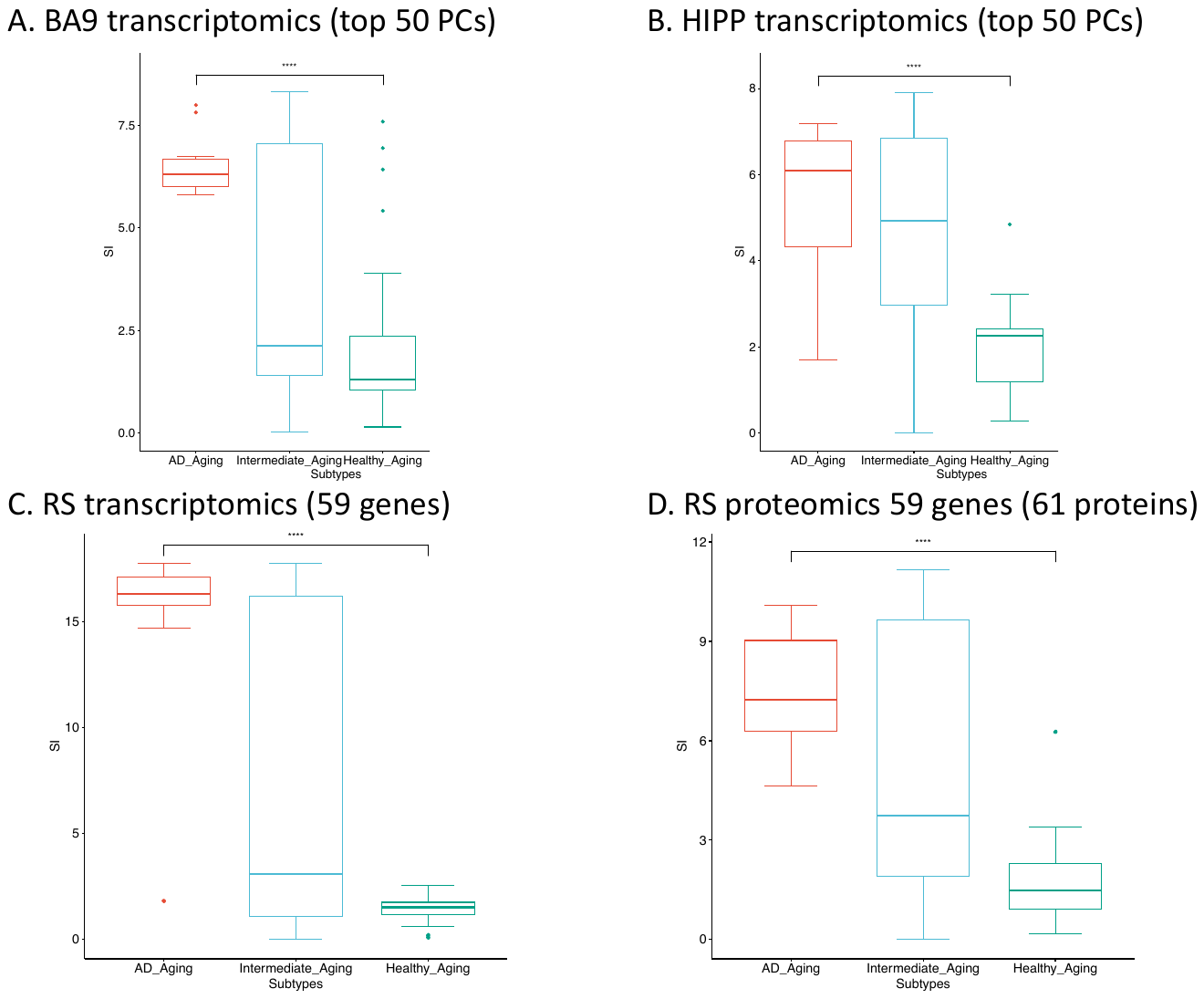

**Figure S12.** 10 CR genes/proteins expression boxplot in GTEx and RS 4 datasets. **A.** 10 CR genes in HIPP GTEx transcriptomics. **B.** 10 CR genes in GTEx PFC transcriptomics. **C.** 10 CR genes in PFC RS proteomics. **D.** 10 CR genes in PFC RS transcriptomics. N for NL, I for INM, A for ADL. Kruskal-Wallis rank sum test (KW) and wilcox.test rank sum test were used to calculate the significance levels between the groups. Wilcox.test: "****", "***", "**", "*", "" for P at 0, 0.0001, 0.001, 0.01, 0.05 and 1 respectively. KW test: P value in top left of each sub-figure.

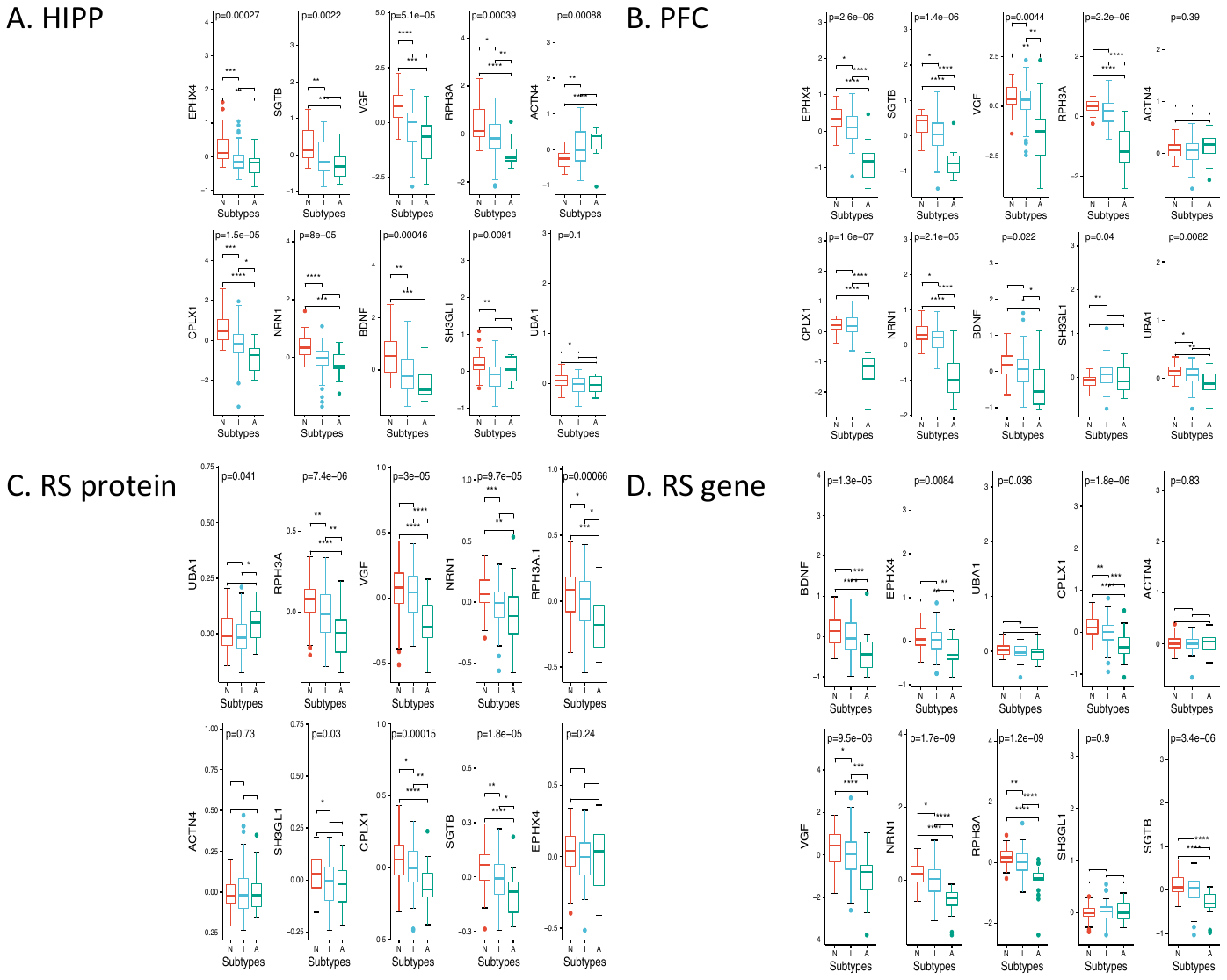

**Figure S13.** GTEx CRBL subgroups and subgroups DEGs vs AD signatures. **A.** GTEx CRBL shows two major subtypes based on 5 subgroups. Aging_SubType: GTEx CRBL NL, ADL, INM subtype. **B.** CRBL pair-wised subgroup DEGs (|Fold change| > 2, FDR < 0.05) overlapped with AD signatures. AD order in DEGs: D, C > E > A, B. **C.** GTEx CRBL, PFC, HIPP subgroups.

**Figure S14.** Top PFC specific ADL vs. NL DEGs (FDR ≤ 0.01) average expression after adjusted covariances. A is for ADL subtype, I is for INM subtype, N is for NL subtype, F is for PFC region, H is for HIPP region. E.g., A_F is for ADL subtype in PFC region. UP.NL or DN.NL is for BaseNL genes due to NL samples gene expression baseline levels. UP.ADL or DN.ADL is for up or down-regulated BaseADL genes due to ADL samples gene expression baseline levels. UP.ALL is for up-regulated gene due to both NL and ADL samples gene expression baseline levels. Kruskal-Wallis rank sum test (KW) and wilcox.test rank sum test were used to calculate the significance levels between the groups. Wilcox.test: "****", "***", "**", "*", "" for P at 0, 0.0001, 0.001, 0.01, 0.05 and 1 respectively.

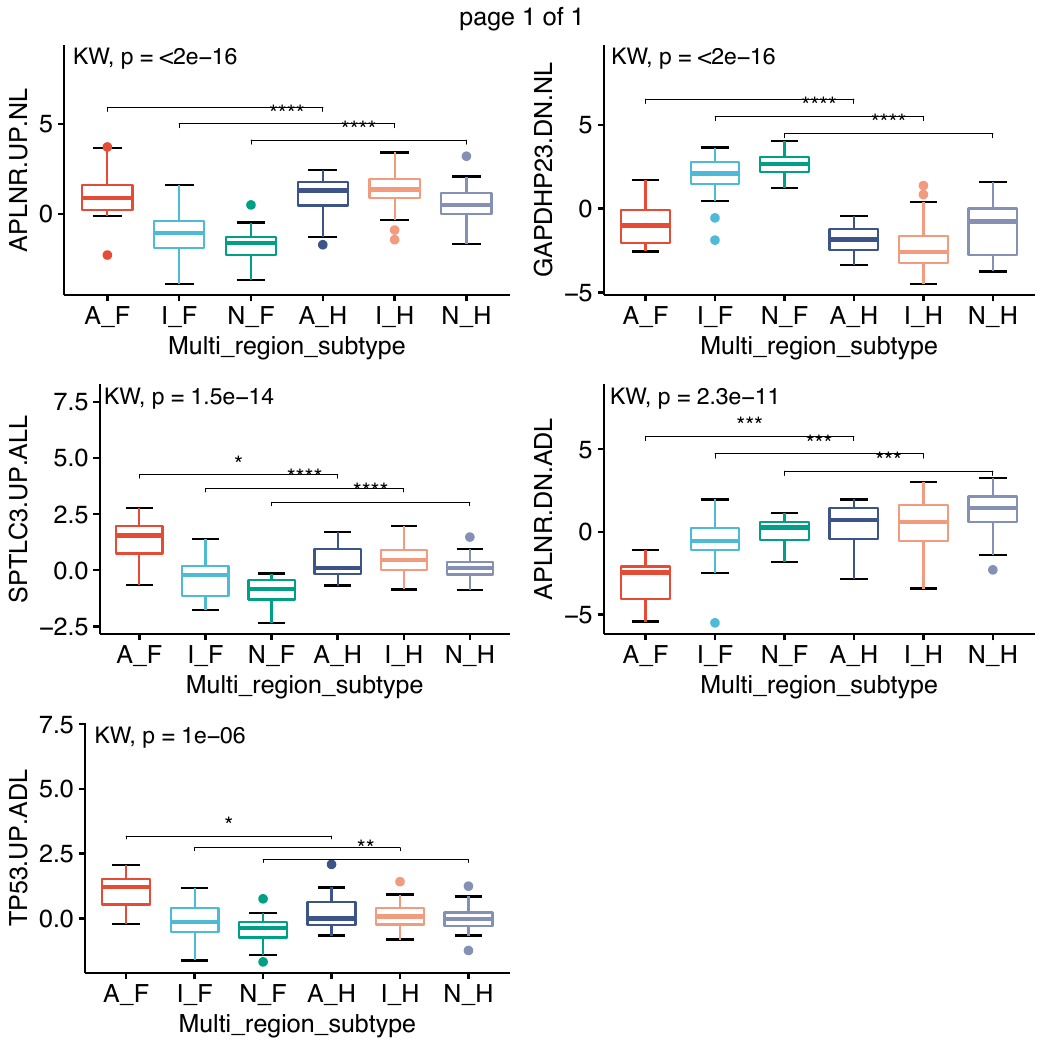

**Tables:**

**Table S1. Description of datasets and aging and AD gene signatures and function annotation of global signatures.**

**Table S1A. List of datasets used for obtaining aging and AD gene signatures.**

| **Datasets** | **Gene List ID** | **List Details (Tissue, traits)** | **# of genes (UP/DOWN)** | **# of protein coding genes** | **Sample Size (AD/N)** |
| --- | --- | --- | --- | --- | --- |
| Transcriptomic “AD” sets (T_AD) | | | | | |
| ROSMAP | ROSMAP | DLPFC | 162(89/73) | 134(76/58) | 241 |
| Jager | Jager_Clinical_AD | Clinical_AD in DLPFC | 855(466/389) | 735(412/323) | 478 |
| Jager | Jager_Cognitive_decline | Cognitive_decline in DLPFC | 3035(1481/1554) | 2627(1258/1369) | 478 |
| Jager | Jager_Tau_tangles | Tau_tangles in DLPFC | 238(155/83) | 207(131/76) | 478 |
| Jager | Jager_B_amyloid | B_amyloid in DLPFC | 2315(1158/1157) | 1998(1056/942) | 478 |
| Jager | Jager_Patho_AD | Patho_AD in DLPFC | 98(58/40) | 85(51/34) | 478 |
| HBTRC | HBTRC | BA9 |  | 1224(637/587) | 549(376/173) |
| Canchi2019 | Canchi2019 | BA9 | 1722 | 1559(665/884) | 414(263/151) |
| AMP AD-PHG | AMPAD_PHG | PHG |  | 1471(777/694) | 215 |
| Annese2018 | Annese2018_HIPP | HIPP | 2122(808/1314) | 1925(742/1183) | 10 |
| Rooij2019 | Rooij2019_HIPP | HIPP | 2840(1109/1731) | 2667(1045/1622) | 28 |
| Proteomic cognitive stability gene sets (P_Aging) | | | | | |
| Wingo2019 | Wingo2019_meta | Meta of BLSA and Banner | 579(350/229) | 538((324/214) | 143(44/99) |
| Wingo2019 | Wingo2019_BLSA | BA9 in BLSA cohorts | 127 | 108(90/18) | 39(16/23) |
| Wingo2019 | Wingo2019_Banner | BA9 in Banner cohorts | 354 | 329(187/142) | 104(28/76) |
| Proteomic “AD” sets (P_AD) | | | | | |
| Seyfried2017 | Seyfried2017_BA9_ADvsNL | BA9 in BLSA cohorts | 463(153/310) | 441(142/299) | 35(20/15) |
| Seyfried2017 | Seyfried2017_BA9_ADvsAsym | BA9 in BLSA cohorts | 237(115/122) | 224(111/113) | 35(20/15) |
| Seyfried2017 | Seyfried2017_BA9_AsymNL | BA9 in BLSA cohorts | 194(48/146) | 180(41/139) | 30(15/15) |
| Ping2020 | Ping2020_BA9_ADvsNL | DLPFC | 2451(1156/1300) | 2333(1107/1231) | 9/10 |
| Ping2020 | Ping2020_BA9_ADvsAsym | DLPFC | 2236(953/1289) | 2135(911/1230) | 9/8 |
| Ping2020 | Ping2020_BA9_AsymNL | DLPFC | 292(170/122) | 280(165/115) | 8/10 |
| Mendonça2019 | Mendonça2019_BA10_LateNL | BA10 | 121(41/80) | 119(40/79) | 10/7 |
| Mendonça2019 | Mendonça2019_BA10_EarlyNL | BA10 | 70(25/45) | 65(23/42) | 9/7 |

BA:Brodmann areas, e.g. BA9, BA10; DLPFC: Dorsolateral prefrontal cortex.; Banner : Banner Sun Health Research Institute cohorts; BLSA: Baltimore Longitudinal Study of Aging cohorts; PHG: parahippocampal gyrus; HIPP: Hippocampus.

**Table S1B. Protein coding gene list of aging and AD signatures.**

**Table S1C. Summary statistics of GTEx and RS data.**

|  | **GTEx BA9** | **GTEx HIPP** | **RS** |
| --- | --- | --- | --- |
| **Sample size** | **129** | **129** | **167** |
| **Sex (Male/Female)** | **99/30** | **97/32** | **69/98** |
| **Age (mean ± s.d.)** | **58.0 ± 10.4** | **57.5 ± 10.9** | **83.5 ± 9.8** |
| **Age (min, max)** | **20 - 70** | **20 - 70** | **64 - 101** |
| **PMI (mean hours ± s.d.)** | **13.7 ± 4.1** | **13.9 ± 4.3** | **11.9 ± 9.2** |
| **RIN (mean ± s.d.)** | **7.3 ± 0.9** | **6.7 ± 0.8** | **5.0 ± 1.5** |
| **Race (White, %)** | **81 (62.8%)** | **87 (67.4%)** | **147 (88.0%)** |

**Table S1D.** The function annotation of global reverse aging regulated genes (NL_UP, NL_DN).

**Table S1E.** The function annotation of global transcriptomic and proteomic AD signatures (TP_AD).

**Table S2.** The best number of subgroups for each dataset by 2 validation criteria* type (test clusters from 2 to 10).

|  | **Rank1** | **Rank2** | **Rank3** | **Rank4** |
| --- | --- | --- | --- | --- |
| **PFC top 5000 gene (top 8000 gene)** | | | | |
| **APN** | kmeans-2 | kmeans-3 (5) | kmeans-5 (H7) | kmeans-6 (H8) |
| **ADM** | kmeans-2 | kmeans-3 (5) | kmeans-5 (7) | kmeans-4 (8) |
| **Connectivity** | hierarchical-2 | hierarchical-3 | hierarchical-4 | hierarchical-5 |
| **Dunn** | hierarchical-8 (3) | hierarchical-9 (4) | hierarchical-10 (5) | kmeans-7 (H6) |
| **Silhouette** | hierarchical-2 | hierarchical-3 | hierarchical-4 | hierarchical-5 |
| **HIPP top 5000 gene^†^** | | | | |
| **APN** | hierarchical-2 | hierarchical-3 | hierarchical-4 | hierarchical-5 |
| **ADM** | hierarchical-2 | hierarchical-3 | hierarchical-4 | hierarchical-5 |
| **Connectivity** | hierarchical-2 | hierarchical-3 | hierarchical-4 | kmeans-2 |
| **Dunn** | hierarchical-2 | hierarchical-3 | kmeans-9 | kmeans-10 |
| **Silhouette** | hierarchical-2 | hierarchical-3 | kmeans-2 | kmeans-3 |
| **RS top 5000 gene (top 8000 gene)** | | | | |
| **APN** | hierarchical-2 | kmeans-2 | hierarchical-3 | hierarchical-6 (3) |
| **ADM** | hierarchical-2 | kmeans-2 | hierarchical-3 | hierarchical-10 (3) |
| **Connectivity** | hierarchical-2 | hierarchical-3 | hierarchical-4 | hierarchical-5 |
| **Dunn** | hierarchical-2 | hierarchical-7 (3) | hierarchical-8 (5) | hierarchical-9 (6) |
| **Silhouette** | hierarchical-2 | hierarchical-3 | hierarchical-4 | hierarchical-5 |
| **RS top 5000 protein (top 8000 gene)** | | | | |
| **APN** | hierarchical-3 (2) | hierarchical-4 (3) | hierarchical-5 (4) | hierarchical-6 (K5) |
| **ADM** | hierarchical-3 (2) | hierarchical-2 (3) | hierarchical-4 | hierarchical-5 |
| **Connectivity** | hierarchical-2 | hierarchical-3 | hierarchical-4 | hierarchical-5 |
| **Dunn** | hierarchical-6 | hierarchical-7 | hierarchical-8 | hierarchical-9 |
| **Silhouette** | hierarchical-2 | hierarchical-3 | hierarchical-4 | kmeans-2 |

*“internal” criteria with connectivity, Dunn’s index, and average Silhouette algorithm; and “stability” criteria with APN (the average proportion of non-overlap), ADM (the average distance between means) algorithm. If the top 8000 gene results changed, we marked in bracket: H for hierarchical cluster; K for k-means clusters. **^†^**We are unable to include the HIPP top 8000 gene results as a result of requiring more than the max server time.

**Table S3.** The function annotation of GTEx PFC and HIPP 3 pair-wise subtype DEGs (ADL vs. NL, ADL vs. INM, INM vs. NL).

**Table S4.** 167 overlapped RS samples in PFC region with consistent NL (green, N = 61) and ADL (red, N = 21) subtype. DEPs AD order: A (ADL-1), C (ADL-2), B (ADL-3) >> E (NL-2), D (NL-1); DEGs AD order: C (ADL-1), D (ADL-2) > E (INM-1) >> A (NL-2) > B (NL-1).

| Multi-Omics | T_C (N=18) | T_D (N=23) | T_E (N=19) | T_A (N=50) | T_B (N=57) |
| --- | --- | --- | --- | --- | --- |
| P_A (N=7) | 1 | 2 | 1 | 1 | 2 |
| P_C (N=22) | 4 | 3 | 1 | 6 | 8 |
| P_B (N=42) | 6 | 5 | 2 | 11 | 18 |
| P_E (N=69) | 5 | 6 | 8 | 25 | 25 |
| P_D (N=27) | 2 | 7 | 7 | 7 | 4 |

Notes:T for transcriptomics, P for proteomics.

**Table S5.** The function annotation of GTEx PFC and RS 3 pair-wise subtype DEGs (ADL vs. NL, ADL vs. INM, INM vs. NL).

**Table S6.** The function annotation of consistent direction genes in GTEx and RS data.

**Table S7.** The function annotation of 235 RS significant correlated genes.

**Table S8.** Comparison of 167 overlapped RS samples in transcriptomics data derived subtypes and proteomics data derived subtypes based on consistent and significant mRNA-protein correlated genes (Num. genes = 59): The ADL subtypes (N = 39) are highlighted in red color and NL subtypes (N = 41) are highlighted in green color. RS transcriptomics AD order: B (ADL-1) >> D (ADL-2) > E (ADL-3) > C (INM-1) > A (NL-1) and RS proteomics AD order as C (ADL-1) > D (ADL-2) > E (INM-1) > B (NL-2), A (NL-1).

| Multi-Omics | T_B (N=16) | T_D (N=37) | T_E (N=24) | T_C (N=31) | T_A (N=59) |
| --- | --- | --- | --- | --- | --- |
| P_C (N=20) | 5 | 4 | 7 | 0 | 4 |
| P_D (N=29) | 7 | 9 | 7 | 4 | 2 |
| P_E (N=31) | 1 | 9 | 2 | 7 | 12 |
| P_B (N=76) | 3 | 15 | 8 | 20 | 30 |
| P_A (N=11) | 0 | 0 | 0 | 0 | 11 |

Notes: T is for transcriptomics and P is for proteomics.

**Table S9.** 80 CRBL subgroups samples overlapped with HIPP- PFC NL and ADL samples.

|  | HIPP-PFC ADL (6) | HIPP-PFC INM (47) | HIPP-PFC NL (27) |
| --- | --- | --- | --- |
| **CRBL B (NL-1, 21)** | **0** | **13** | **8** |
| **CRBL A (NL-2, 29)** | **1** | **14** | **14** |
| **CRBL C (ADL-1, 5)** | **1** | **2** | **2** |
| **CRBL D (ADL-2, 7)** | **1** | **5** | **1** |
| **CRBL E (INM-1, 18)** | **3** | **13** | **2** |

Notes: **The number of samples for** **CRBL-HIPP-PFC NL: 22, ADL:2, INM: 56.**

**Table S10.** The function annotation of CRBL ADL vs. NL DEGs.

**Table S11.** A summary list of numbers of DEGs between 3 subtypes derived from PFC-HIPP multi-region and CRBL data.

|  | ADL vs. NL | ADL vs. INM | INM vs. NL |
| --- | --- | --- | --- |
| PFC-HIPP Multi-region: HIPP DEGs with fold change at 2, FDR ≤ 0.01 | | | |
| Down | 1366 | 0 | 528 |
| Up | 721 | 0 | 221 |
| PFC-HIPP Multi-region: PFC DEGs with fold change at 2, FDR ≤ 0.01 | | | |
| Down | 2872 | 2293 | 0 |
| Up | 3128 | 2170 | 0 |
| CRBL region: CRBL DEGs with fold change at 2, FDR ≤ 0.01 | | | |
| Down | 196 | 391 | 94 |
| Up | 640 | 335 | 372 |

Notes: The number of samples used for DEG calculation- PFC-HIPP:12 ADL, 28 NL, 68 INM; CRBL: 24 ADL, 113 NL, 28 INM.

**Table S12.** Geometric mean expression of four representative pathway genes in NL and ADL samples across CRBL, PFC, and HIPP.

|  | Inflammatory Response  GO:0006954  (699 genes)* | extracellular exosome GO:0070062  (83 genes) | synapse GO:0045202 (1524 genes) | chemical synaptic transmission GO:0007268 (743 genes) |
| --- | --- | --- | --- | --- |
| HIPP NL mean | 1.067 | 1.031 | 1.09 | 1.107 |
| PFC NL mean | 0.992 | 0.928 | 1.148 | 1.234 |
| CRBL NL mean | 0.874 | 0.925 | 0.932 | 0.926 |
| HIPP ADL mean | 1.216 | 1.176 | 0.902 | 0.835 |
| PFC ADL mean | 1.277 | 1.182 | 0.944 | 0.901 |
| CRBL ADL mean | 0.96 | 1.02 | 0.933 | 0.909 |
| HIPP ADL vs. NL (%^†^) | 0.139 | 0.140 | - 0.173 | - 0.245 |
| PFC ADL vs. NL (%) | 0.288 | 0.274 | - 0.178 | - 0.270 |
| CRBL ADL vs. NL (%) | 0.098 | 0.103 | - 0.001 | - 0.018 |

***the number of genes detected in our expression data. ^†^change percent (%) = (ADL_mean – NL_mean)/NL_mean.**

**Table S13.** A summary list of numbers of DEGs or DEPs between subgroups derived from multi-region and multi-omics data.

|  | ADL vs. NL | ADL vs. INM | INM vs. NL |
| --- | --- | --- | --- |
| GTEx Multi-region:HIPP DEGs with FDR ≤ 0.01 | | | |
| Down | 2042 | 0 | 2448 |
| Up | 1298 | 0 | 1709 |
| GTEx Multi-region:PFC DEGs with FDR ≤ 0.01 | | | |
| Down | 4424 | 3628 | 0 |
| Up | 4030 | 4144 | 0 |
| RS Multi-Omics: PFC DEGs with FDR ≤ 0.01 | | | |
| Down | 4210 | 2583 | 0 |
| Up | 2100 | 913 | 0 |
| RS Multi-Omics: PFC DEPs with FDR ≤ 0.01 | | | |
| Down | 600 | 14 | 88 |
| Up | 665 | 4 | 152 |

Notes: The number of samples used for DEG/DEP calculation- GTEx:12 ADL, 28 NL, 68 INM; RS: 21 ADL, 61 NL, 85 INM.

**Table S14.** PFC specific ADL vs. NL DEGs (FDR ≤ 0.01) can be divided into several group including BaseNL (different NL gene expression, no significant difference in ADL samples between PFC and HIPP), BaseADL (different ADL gene expression but no significant difference in NL samples between PFC and HIPP), BaseALL (different gene expression in both NL and ADL samples), and Others (all other situations). The number of significant DEGs are listed in the table.

| DEGs counts | UP (1148) | DOWN (1080) |
| --- | --- | --- |
| BaseNL | 330 | 789 |
| BaseALL (NL, ADL) | 6 | 0 |
| BaseADL | 167 | 4 |
| Others | 645 | 287 |

**Table S15.** The function annotation of 330 up-regulated and 789 down-regulated BaseNL genes.

**Table S16.** Glial and Endothelial cell type proportion in AD-like subgroups.

| Tissue | AD-like order groups | Ast | Oli | Mic | End |
| --- | --- | --- | --- | --- | --- |
| GTEx HIPP | A, B > C | B | A | B > C | A, B |
| GTEx BA9 | E > D, B | B | E | D | B, D |

Notes: astrocytes (Ast), endothelial cells (End), microglia (Mic), oligodendrocytes (Oli).
